# Adipocyte autophagy limits gut inflammation by controlling oxylipin levels

**DOI:** 10.1101/2021.10.25.465200

**Authors:** Felix Clemens Richter, Matthias Friedrich, Nadja Kampschulte, Mathilde Pohin, Ghada Alsaleh, Irina Guschina, Sarah Karin Wideman, Errin Johnson, Mariana Borsa, Klara Piletic, Paula Hahn, Henk Simon Schipper, Claire M. Edwards, Rudolf Zechner, Nils Helge Schebb, Fiona Powrie, Anna Katharina Simon

## Abstract

Lipids play a major role in inflammatory diseases by altering inflammatory cell functions, through their use as energy substrates or as lipid mediators such as oxylipins. Autophagy, a lysosomal degradation pathway that limits inflammation, is known to impact on lipid availability, however whether this controls inflammation remains unexplored. We found that upon intestinal inflammation visceral adipocytes upregulate autophagy and that adipocyte-specific loss of the autophagy gene *Atg7* exacerbates inflammation. While autophagy decreased lipolytic release of free fatty acids, loss of the major lipolytic enzyme *Pnpla2/Atgl* in adipocytes did not alter intestinal inflammation, ruling out free fatty acids as anti-inflammatory energy substrates. Instead, *Atg7*-deficient adipose tissues exhibited an altered oxylipin balance, driven through an NRF2-mediated upregulation of *Ephx1*. This was accompanied by a shift in adipose tissue macrophage polarization with reduced secretion of IL-10, leading to lower circulating levels of IL-10. These results suggest an underappreciated fat-gut crosstalk through an autophagy-dependent regulation of anti-inflammatory oxylipins, indicating a protective effect of adipose tissues for distant inflammation.

## Introduction

Autophagy is an essential cellular recycling pathway that engulfs cellular contents, including organelles and macromolecules, in a double membraned autophagosome and directs them towards lysosomal degradation. Many cell types, including immune cells, are reliant on autophagy during their differentiation and for their functions^1^. Consequently, autophagy dysfunction is associated with the development of a variety of inflammatory diseases and metabolic disorders^2, 3^. Inflammatory bowel diseases (IBD) including its two predominant manifestations, Crohn’s disease (CD) and ulcerative colitis (UC), describe a complex spectrum of intestinal inflammation. Genome wide association studies identified autophagy-related genes as susceptibility alleles in IBD^4–6^. Mechanistic studies revealed that ablation of autophagy in immune and epithelial cells promotes intestinal inflammation^7–9^. In addition to the strong genetic association of autophagy and IBD, patients with CD often present with an expansion of the mesenteric adipose tissue around the inflamed intestine, indicating an active involvement of the adipose tissue in the disease pathology^10^.

Adipose tissues represent an important immunological organ harbouring a variety of immune cells, which are highly adapted to live in lipid-rich environments, such as adipose tissue macrophages (ATMs)^11^. Lean adipose tissues are predominantly populated by tissue-resident M2-type ATMs, while inflammation, such as induced by obesity, subverts their homeostatic functions and promotes pro-inflammatory M1-type polarization^12^. Polarization and function of ATMs depend on the integration of a variety of inflammatory and metabolic signals. M2-type macrophages rely on the availability and uptake of lipids and the subsequent metabolization of free fatty acids (FFA) compared to M1-type macrophages^13^. In addition, oxygenated polyunsaturated fatty acids, so called oxylipins, which are produced through enzymatic lipid oxidation can be released from adipocytes to modify macrophage cytokine production^14^. Oxylipins have been widely described as regulatory lipid mediators that regulate inflammatory processes and resolution^15–17^. It is plausible that both availability of energy substrates such as FFA and signalling through oxylipin mediators will modulate immune responses. Autophagy contributes to FFA release^18^ and lipid peroxidation^19^, however, to-date, little is known about the impact of autophagy in adipocytes on these metabolic cues and whether these may affect inflammation.

Here, we sought to investigate the impact of adipocyte autophagy on the immune system during inflammation of a distant organ, the intestine. We observed that autophagy is induced in mature adipocytes upon dextran sulphate sodium (DSS)-induced intestinal inflammation, and that loss of autophagy in adipocytes exacerbated gut inflammation. Mechanistically, while autophagy in mature adipocytes is required for the optimal release of FFA during inflammation, this was not causative for increased intestinal inflammation. Instead, loss of adipocyte autophagy stabilized the oxidative stress master transcription factor NRF2 and promoted the oxylipin pathway activity shifting the balance of intra-tissual oxylipins. Local oxylipin dysbalance limited the production of anti-inflammatory IL-10 from ATMs, aggravating intestinal inflammation. Taken together, we demonstrate a novel mechanism of autophagy in adipocytes regulating local oxylipins that promote the anti-inflammatory fat-gut crosstalk, highlighting the importance of inter-tissual control of inflammation.

## Results

### DSS-induced intestinal inflammation promotes autophagy in adipose tissues

To investigate whether autophagy in mature adipocytes is altered in response to intestinal inflammation, we deployed a mouse model of intestinal inflammation evoked by the administration of DSS in drinking water (Figure 1A). As expected, treatment with DSS led to an increased histopathological inflammation, shortened colon length, enlarged mesenteric lymph nodes, and elevated circulating levels of the pro- inflammatory cytokine TNFα (Figure S1A-D). In addition, DSS treatment resulted in a significantly higher infiltration of immune cells in the inflamed colon, predominantly of myeloid origin (Figure S1E- F). Furthermore, DSS colitis reduced body weight (Figure 1B), and in line with that, visceral adipose tissue mass (Figure 1C), as well as serum FFA levels (Figure 1D).

**Figure 1:**
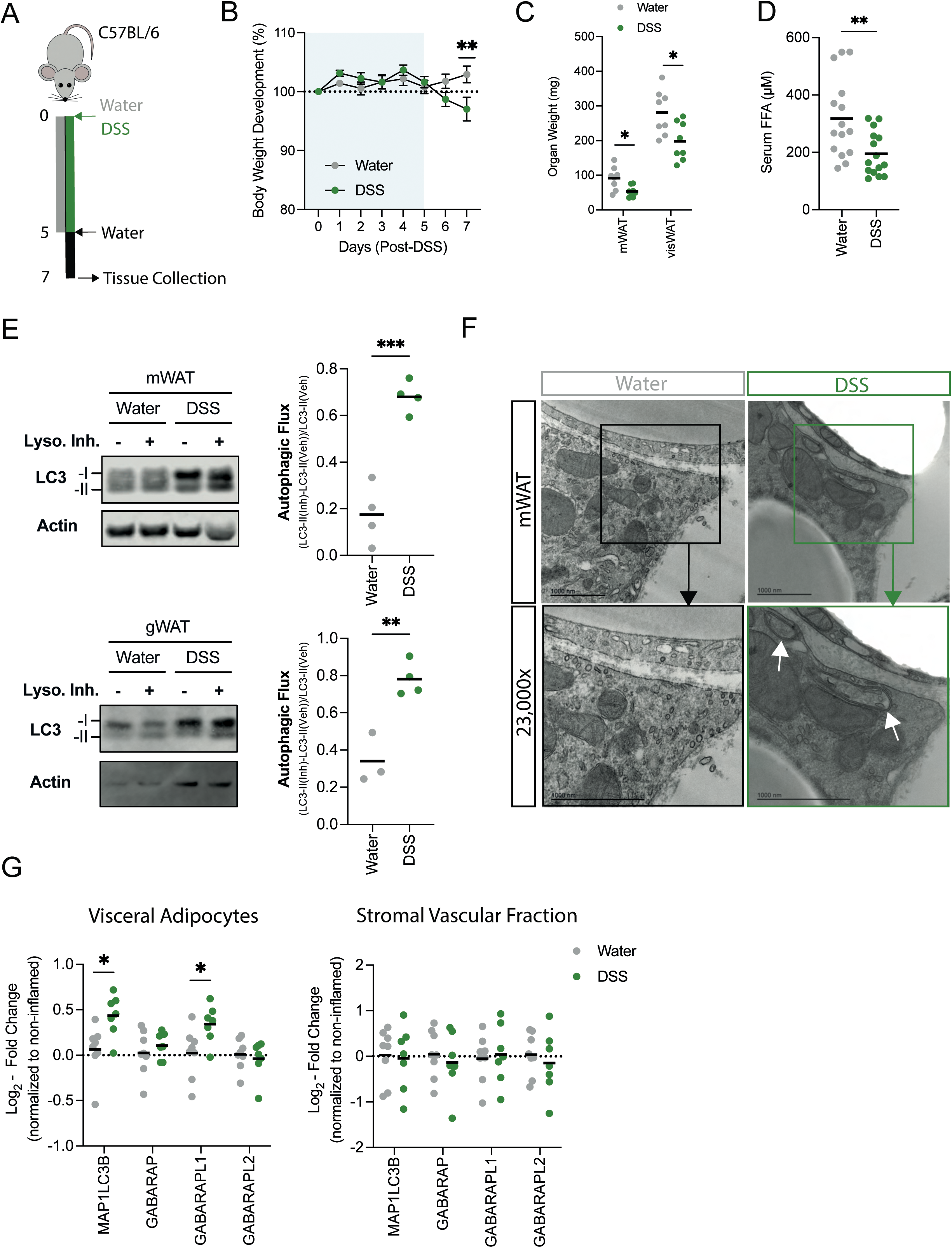
DSS-induced intestinal inflammation promotes autophagy in adipose tissues. (A) Schematic of experimental design. Sex-matched and age-matched wild-type mice were treated for 5 days with 1.5-2% DSS in drinking water, before switched to water for two more days. Mice were sacrificed at day 7 post-DSS induction. (B) Body weight development upon DSS treatment (n = 10-11/group). (C) Tissue weights measured in mesenteric (mWAT) and collective visceral white adipose tissue (visWAT), consisting of gonadal (gWAT), retroperitoneal and omental white adipose tissue at day 7 after start of DSS regime (n = 7-8/group). (D) Circulating serum levels of FFA during DSS-induced colitis at day 7 (n = 15/group). (E) Immunoblot analysis of autophagic flux in mWAT (upper panel) and gWAT (lower panel) adipose tissue stimulated *ex vivo* with lysosomal inhibitors 100nM Bafilomycin A1 and 20mM NH4Cl for 4 hours or DMSO (Vehicle) (n = 3-4/group). (F) Representative transmission electron microscopy images from mesenteric adipose tissue 7 days post DSS-induced colitis induction. Lower panel is showing magnification of selected area. White arrows show autophagosomal structures. (G) *Atg8* homologues expression was measured by qPCR in visceral adipocytes fraction (right) and stromal vascular fraction (left panel) during DSS-induced colitis (n = 7-8/group). Data are represented as mean. (B) Two-Way ANOVA. (C.D,E,G) Unpaired Student’s t-test.

To assess changes in autophagy levels, adipose tissue explants from water- or DSS-treated animals were cultured in the absence or presence of lysosomal inhibitors and the accumulation of the lipidated autophagosomal marker LC3 protein (LC3-II) was quantified. DSS-induced intestinal inflammation substantially increased autophagic flux in mesenteric and in gonadal white adipose tissue (mWAT and gWAT, respectively) (Figure 1E), indicating that both adipose tissues proximal and distal to the intestine are responsive to the inflammation. To validate that adipocytes, but not other adipose tissue-resident cell types, contribute to the increased autophagic flux in the adipose tissue, we first prepared adipose tissue for transmission electron microscopy. Autophagosomal double membrane structures were readily identified in adipocytes from DSS-treated mice (Figure 1F). Additionally, enriched adipocytes fractions increased the expression of several *Atg8* homologues upon DSS colitis, further demonstrating an induction of autophagy in this cell type (Figure 1G). In contrast, the adipocyte-depleted stromal vascular fraction containing other adipose tissue-resident immune and stromal cells showed no transcriptional changes in *Atg8* expression (Figure 1G). Overall, these results demonstrate that autophagy is induced in adipocytes in response to DSS-induced intestinal inflammation.

### Loss of adipocyte autophagy exacerbates intestinal inflammation

Given the increased adipose autophagy we observed as reaction to intestinal inflammation, we next investigated whether loss of autophagy in adipocytes affects intestinal inflammation. To exclude developmental effects of autophagy loss^18, 20^, we used a tamoxifen-inducible knockout mouse model to ablate the essential autophagy gene *Atg7* specifically in mature adipocytes (*Atg7^Ad^*) in adult mice (Figure 2A). Tamoxifen administration led to the significant reduction of *Atg7* transcript levels in visceral adipocytes (Figure 2B). This deletion was further confirmed at the protein level (Figure 2C). Importantly, the adipocyte-specific loss of ATG7 resulted in the interruption of conversion of LC3-I to LC3-II in the adipose tissue (Figure 2C) confirming effective disruption of the autophagic process in adipose tissue. Having confirmed efficient deletion of *Atg7* and disruption of autophagic flux in adipocytes, we next compared the effects of autophagy loss in adipocytes in steady state and DSS-induced colitis (Figure 2D). In all assessed parameters, loss of adipocyte autophagy in steady state mice had no effects on intestinal immune homeostasis. In contrast, *Atg7^Ad^* mice showed an increased loss of body weight in comparison to littermate controls upon DSS-treatment (Figure 2E). In addition, *Atg7^Ad^* mice treated with DSS had significantly shorter colon when compared to their wild-type littermates during acute inflammation (Figure 2F). Blinded histopathological assessment^21^ confirmed that DSS-treated *Atg7^Ad^* mice exhibited more severe tissue damage accompanied by increased inflammation and reduced features of repair throughout the colon (Figure 2G). Consistent with an increased inflammatory response, we found increased gene expression of alarmins such as *Il1a* and *Il33*, pro-inflammatory cytokines *Tnfa*, *Ptx3*, *Ifng* and the IFNγ-regulated chemokine *Cxcl9* in *Atg7^Ad^* mice (Figure 2H). Although total CD45^+^ immune cells numbers were comparable between adipocyte autophagy-deficient mice and littermate controls (Figure 2I), DSS-inflamed *Atg7^Ad^* mice showed an increased frequency of monocytes infiltrating the intestinal tissue (Figure 2J). In particular, the number of MHCII-expressing, inflammatory monocytes were increased in the lamina propria of *Atg7^Ad^* mice (Figure 2K). This phenotype is in line with the fact that autophagic flux was predominantly induced in adipose tissues upon intestinal damage by DSS (Figure 1E) suggesting an important function of adipocyte autophagy during intestinal inflammation. Taken together, these data demonstrate that loss of adipocyte autophagy exacerbates intestinal inflammation in the acute phase of DSS-induced colitis.

**Figure 2:**
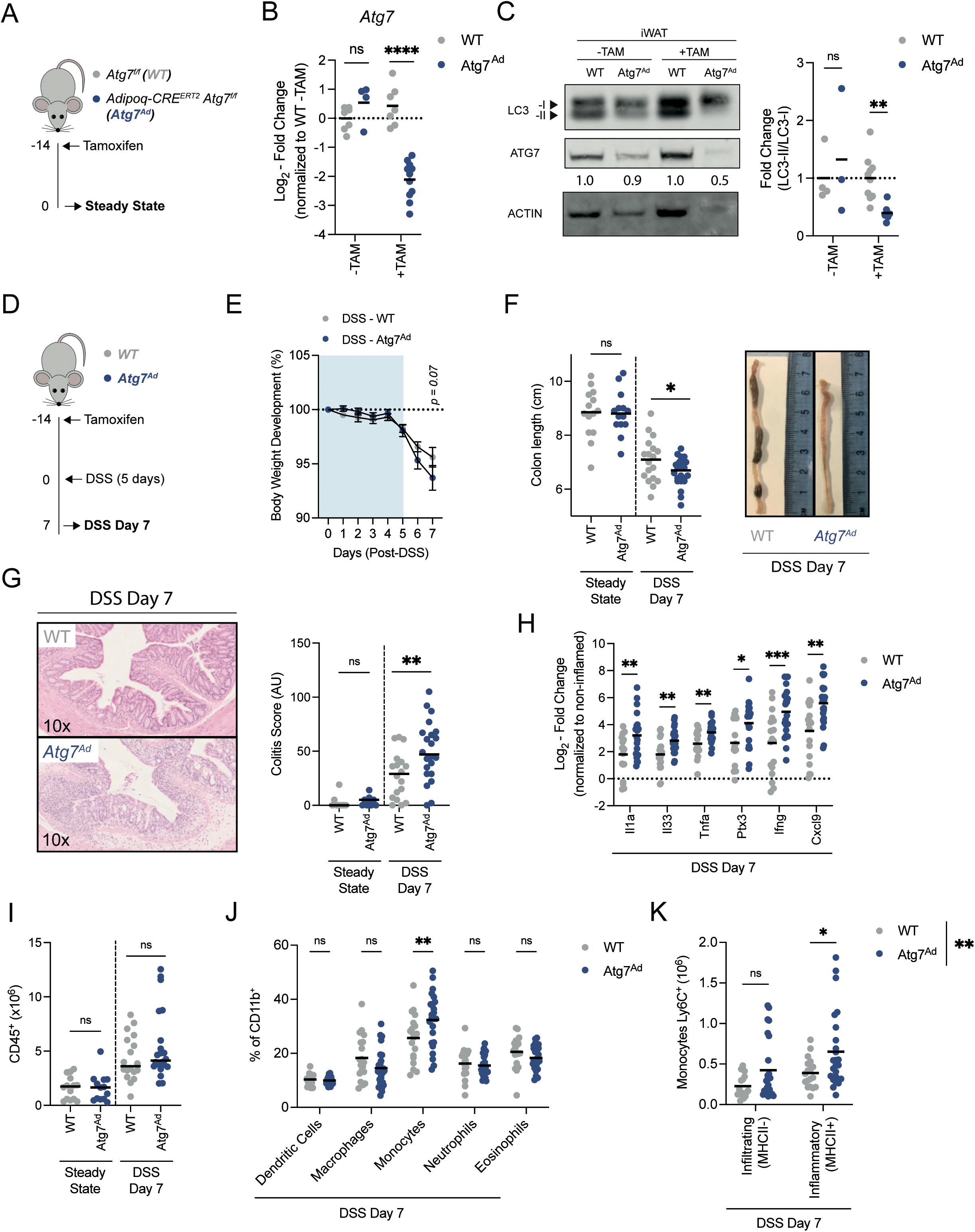
Loss of adipocyte autophagy exacerbates DSS-induced colitis. (A) Schematic of experimental design. Sex-matched and age-matched littermates were treated at 8-12 weeks of age with tamoxifen for five consecutive days before tissues were analysed 14 days after the last tamoxifen administration (Steady State). (B) Representative quantification of knock-out efficiency measured on *Atg7* transcript level by qRT-PCR in purified primary visceral adipocyte at two weeks post-tamoxifen treatment (n = 4-11/group). (C) Representative immunoblot for ATG7 and LC3-I/II protein expression and quantification of LC3 conversion ratio (LC3-II/LC3-I) (n = 3-10/group). (D) Schematic of experimental design. Sex-matched and age-matched littermates were treated at 8-12 weeks of age with tamoxifen for five consecutive days and DSS-induced colitis was induced after a two-week washout phase (DSS Day 7). (E) Body weight development upon DSS treatment; n = 25/group. (F) Colon length after two weeks post-deletion (steady state; n = 14/group) and after DSS at day 7 (n = 18-22/group). (G) Representative H&E staining images (10x magnification) of colon sections and quantification of histological score at steady state (n = 9/group) and DSS colitis (n = 18-22/group). (H) Expression of pro-inflammatory cytokines in colon tissues at 7 days post-DSS induction; n = 18- 22/group pooled from three independent experiments. Dotted line represents uninflamed controls. (I) Absolute number CD45^+^ immune cells from colons at steady state (n = 13-14/group) or at 7 days post-DSS induction (n = 18-22/group). (J) Frequency of myeloid cell population in colon at day 7 post-DSS induction (n = 18-22/group). (K) Absolute number of Ly6C^+^ monocytes discriminated by the absence or presence of MHCII for infiltrating and inflammatory monocytes respectively (n = 18-22/group). Data are represented as mean. (E,J,K) Two-Way ANOVA. (B,C,F,G,H,I) Unpaired Student’s t-test.

Since intestinal inflammation induced by DSS is self-resolving, we assessed the impact of adipocyte autophagy loss during resolution of the inflammation (Figure S2A). Two weeks after initial DSS administration, we did not find any differences in colon length between *Atg7^Ad^* and littermate controls and, equally there were no significant histopathological differences observed between the groups (Figure S2B-C). Interestingly, frequencies and total numbers of colonic FOXP3^+^ regulatory T cells (Tregs) were decreased in adipocyte autophagy-deficient animals compared to wild-type animals (Figure S2D), despite not affecting disease recovery. Intestinal FOXP3^+^ Tregs are classified into three distinct subsets based on co-expression of TH2 and TH17 transcription factors GATA3^+^ and RORgt^+^, respectively^22^. While all populations tended to be diminished in *Atg7^Ad^* mice, only RORgt^-^ FOXP3^+^ Tregs were significantly reduced (Figure S2E). These data suggest that adipocyte autophagy is dispensable for the resolution of DSS-induced inflammation but may affect expansion of intestinal Tregs in response to intestinal tissue injury.

### Intestinal inflammation promotes a lipolytic transcriptional profile in primary adipocytes

At this point, it remained unclear how adipocytes regulate intestinal inflammation. We hypothesized that visceral adipocytes would alter their transcriptional inflammatory profile during intestinal inflammation. To test this, visceral adipocytes were collected from wild-type and *Atg7^Ad^* mice treated with water or DSS and subjected to RNA sequencing. Since we anticipated sex-specific differences in adipocyte transcription profiles, we included the same number of male and female mice in each experimental group. The treatment clearly separated the experimental groups in the principal component analysis (PCA) (Figure 3A). As expected, sex-specific transcriptional changes explained ∼33% of the dataset variance (Figure 3A), in line with previous reports^23^. Next, we compared non-inflamed to inflamed adipocytes by regressing genotype and sex to identify the impact of intestinal inflammation on the adipocyte transcriptome. More than 4700 genes were differentially regulated between these states (Figure 3B), among which 2415 were significantly upregulated and 2333 downregulated. Gene ontology analysis of these differentially expressed genes revealed an enrichment in several pathways (Figure 3C). Confirming our earlier results that adipocyte autophagy is affected by DSS-induced colitis (Figure 1G), intestinal inflammation led to an enrichment of genes involved in macroautophagy in visceral adipocytes (Figure S3A), including an increased expression of several *Atg8* homologues (*Gabarap*, *Gabarapl1*, *Map1lc3a*, *Map1lc3b*) (Figure S3B). In addition, genes related to fatty acid metabolism were enriched in visceral adipocytes during intestinal inflammation (Figure 3D). Interestingly, genes encoding key proteins involved in the lipolytic pathway, *Adrb3*, *Pnpla2*, *Lipe* and *Fabp4,* were upregulated upon intestinal inflammation (Figure 3E), suggesting a change in the lipolytic status of the adipocytes. Similar to cachexic conditions, an increase of lipolytic genes (*Lipe*, *Pnpla2*) and simultaneous decrease of lipogenic genes (*Dgat2*, *Mogat2*, *Lpl*) is observed^24^. We confirmed protein levels of these enzymes and found that DSS-induced colitis increased phosphorylation of hormone-sensitive lipase (HSL) and the expression of ATGL, two key enzymes in the lipolytic pathway, in line with increased lipolytic activity from the adipocytes (Figure 3F). Overall, intestinal inflammation leads to a broad transcriptional response in visceral adipocytes, altering autophagy and fatty acid metabolism, which are reminiscent of a cachexic response phenotype^24^. Based on their main function in lipid provision, we hypothesized that autophagy may post-transcriptionally affect the release of FFA from adipocytes and thus may control nutrient availability to immune cells.

**Figure 3:**
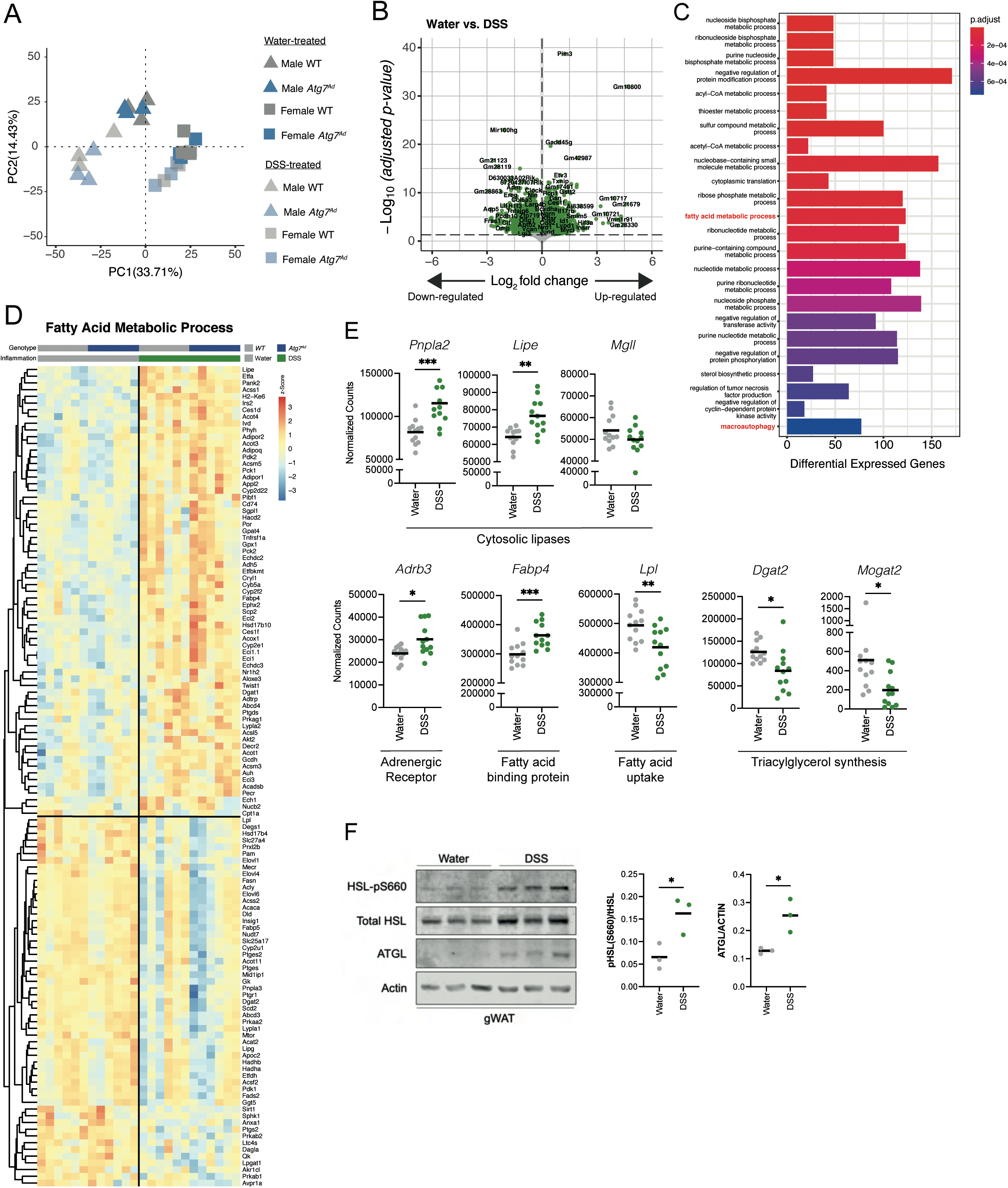
Intestinal inflammation induces fatty acid metabolic transcriptional programs in primary visceral adipocytes. (A) Principal component analysis of all mice revealing a strong sex effect in the overall transcriptome. (B) Differential gene expression assessing transcriptional changes associated with DSS-induced inflammation after regressing effect of sex and genotypes in visceral adipocytes. (C) Pathway enrichment analysis of significantly differentially expressed genes in visceral adipocytes during DSS colitis. (D) Heatmap representing differentially expressed genes associated in fatty acid metabolism during DSS-induced colitis in visceral adipocytes. (E) Normalized counts of selected key enzymes and proteins involved in the lipolysis pathway and lipogenic pathway in visceral adipocytes (n = 12/group). (F) Representative immunoblot for key lipolytic enzymes HSL and ATGL protein expression and quantification (n = 3/group) from one independent experiment. Data are represented as mean. (E,F) Unpaired Student’s t-test.

### Adipocyte autophagy regulates FFA secretion

Recent reports implicated autophagy in mature adipocytes in the secretion of FFA in response to β- adrenergic receptor-mediated lipolysis^19, 25^. To confirm the importance of adipocyte autophagy for optimal lipolytic output, adipose tissue explants were stimulated with the β-adrenergic receptor agonist isoproterenol and FFA levels were quantified. As expected, FFA secretion was reduced upon lipolysis stimulation in autophagy-deficient as compared to autophagy-proficient adipocytes (Figure 4A). TNFα is a crucial cytokine for human and murine IBD pathologies^26^ and can affect adipose tissue through inhibition of lipogenesis and by promoting FFA secretion^27^. Since circulating TNFα levels were elevated during DSS colitis (Figure S1D), we investigated its effects on adipocyte lipid metabolism. Expression of the gene encoding for TNF receptor 1, *Tnfrsf1a*, was upregulated during DSS-induced inflammation in both genotypes, suggesting that TNFα-sensing was unaffected by the loss of adipocyte autophagy (Figure 4B). Next, we assessed the impact TNFα on FFA release from adipose tissue explants from *Atg7^Ad^* mice or littermate controls. In the presence of TNFα, adipocytes increase FFA secretion, however strikingly, this was significantly blunted in autophagy-deficient adipocytes (Figure 4C). Consistent with the decreased lipolytic activity of autophagy-deficient adipocytes, *Atg7^Ad^* mice exhibit reduced serum FFA levels compared to wild-type littermates upon DSS colitis (Figure 4D). While we established that autophagy could modulate overall FFA release, we next tested whether autophagy affects the production and secretion of specific FFA species. To investigate this, serum samples from water and DSS-treated animals were analysed by GC-FID. Confirming our initial findings, the serum concentration of many FFA species was reduced upon adipocyte autophagy loss, indicating that adipocyte autophagy controls overall FFA levels rather than specific FFAs (Figure 4E). Interestingly, loss of adipose tissue mass was comparable between both genotypes upon DSS-induced colitis (Figure S4A).

**Figure 4:**
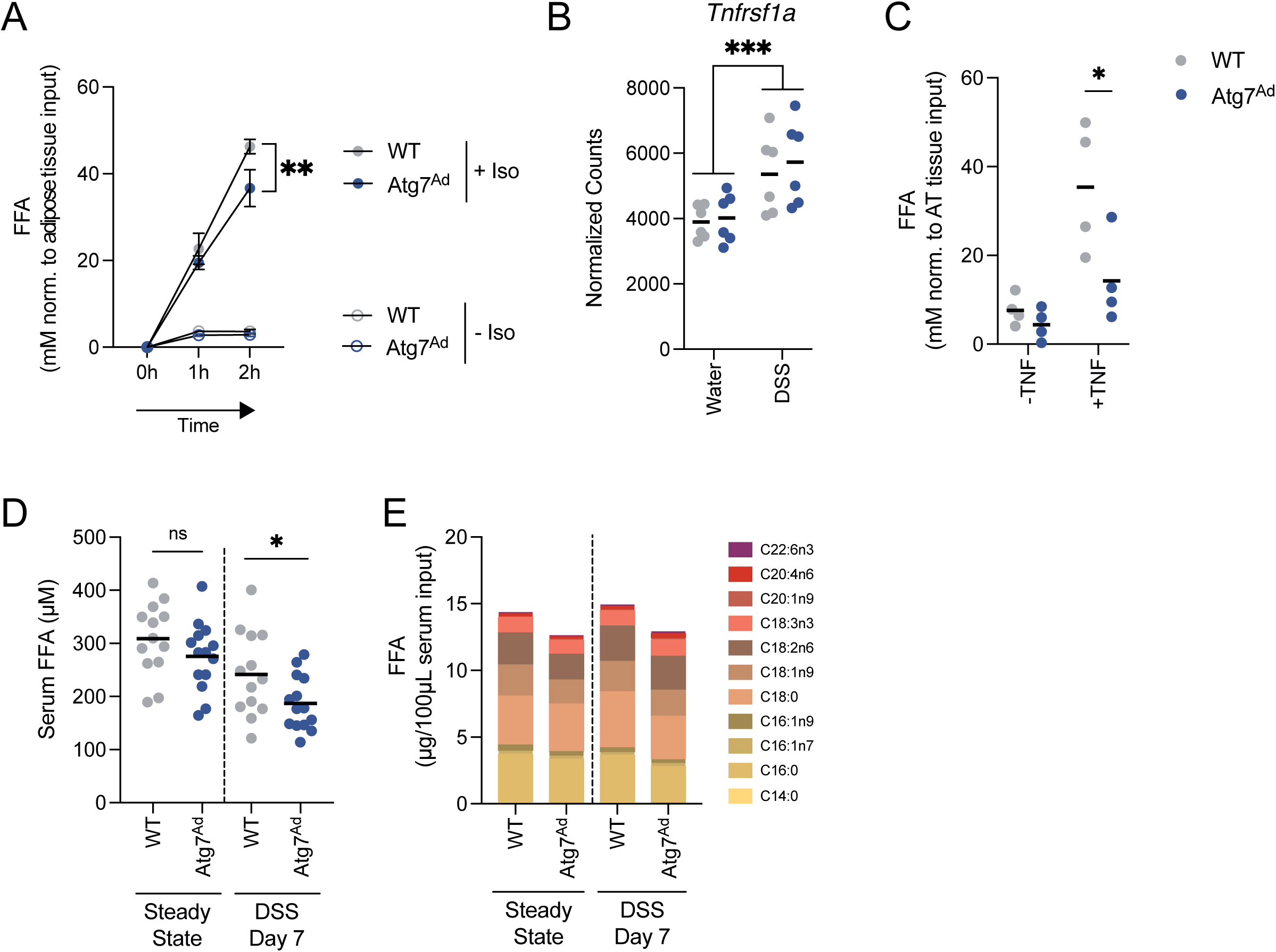
Autophagy loss reduces secretion of fatty acids from adipocytes. (A) *Ex vivo* lipolysis assays on *Atg7*-deficient adipose tissue explants simulated with isoproterenol (10µM) for 1-2h; n = 4-5/group representative for three independent experiments. (B) Expression of the gene encoding TNF receptor 1, *Tnfrsf1a*, on visceral adipocytes during intestinal inflammation. Data expressed as normalized counts from transcriptome analysis. (C) *Ex vivo* lipolysis assay on *Atg7*-deficient adipose tissue explants simulated with TNFα (100ng/mL) for 24h before replacing with fresh medium in the absence of TNFα for 3h (n = 4/group). (D) Serum levels of circulating FFAs measured in wild-type and *Atg7*-deficient mice (n = 13-14/group). (E) Concentration of individual FFA species in serum in water-treated and DSS-treated mice as measured by FID-GC (n = 12-14/group). Data are represented as mean. (A-C,E) Two-Way ANOVA. (D) Unpaired Student’s t-test.

It has previously been described that the adipokines leptin and adiponectin can influence intestinal inflammation in both pre-clinical and clinical situations^28, 29^, we therefore assessed the impact of adipocyte autophagy loss on circulating levels of these adipokines. The levels of both adipokines were equally reduced in their circulation, paralleling the general loss of adipose tissue mass (Figure S4B-C). Taken together, our data suggests that adipocyte autophagy blunts the release of FFA both upon β- adrenergic receptor- and TNFα-mediated lipolysis.

### Adipocyte lipolysis is dispensable for DSS-induced colitis severity

Based on our data, we hypothesized that differences in FFA availability may be responsible for a differential intestinal immune response. We therefore sought to determine the importance of adipocyte lipolysis during DSS-induced colitis (Figure 5A) by deleting the cytoplasmic lipase ATGL, a rate-limiting enzyme in the lipolytic pathway^30^. Using adipocyte-specific *Pnpla2/Atgl* (*Atgl^Ad^*) knockout mice, we first confirmed that *Pnpla2/Atgl* was efficiently deleted in purified visceral adipocytes (Figure 5B). Strikingly, *Atgl^Ad^* mice lost comparable amounts of body weight upon DSS-induced colitis (Figure 5C), although adipose tissue loss was completely prevented (Figure 5D). This data underlines that *Atgl*-driven lipolysis is a main driver for adipose tissue loss during DSS-induced colitis. Detailed analysis of the colon showed no changes in colon shortening, histopathological scores and expression of inflammatory cytokines (Figure 5E-G). Similarly, there was no difference in the infiltration and presence of different pro-inflammatory immune cell population in the colonic lamina propria (Figure 5G-J). In summary, adipocyte lipolysis does not affect acute intestinal inflammation, suggesting that provision of FFA is unlikely to be the mechanism by which autophagy in adipocytes exerts its anti-inflammatory role.

**Figure 5:**
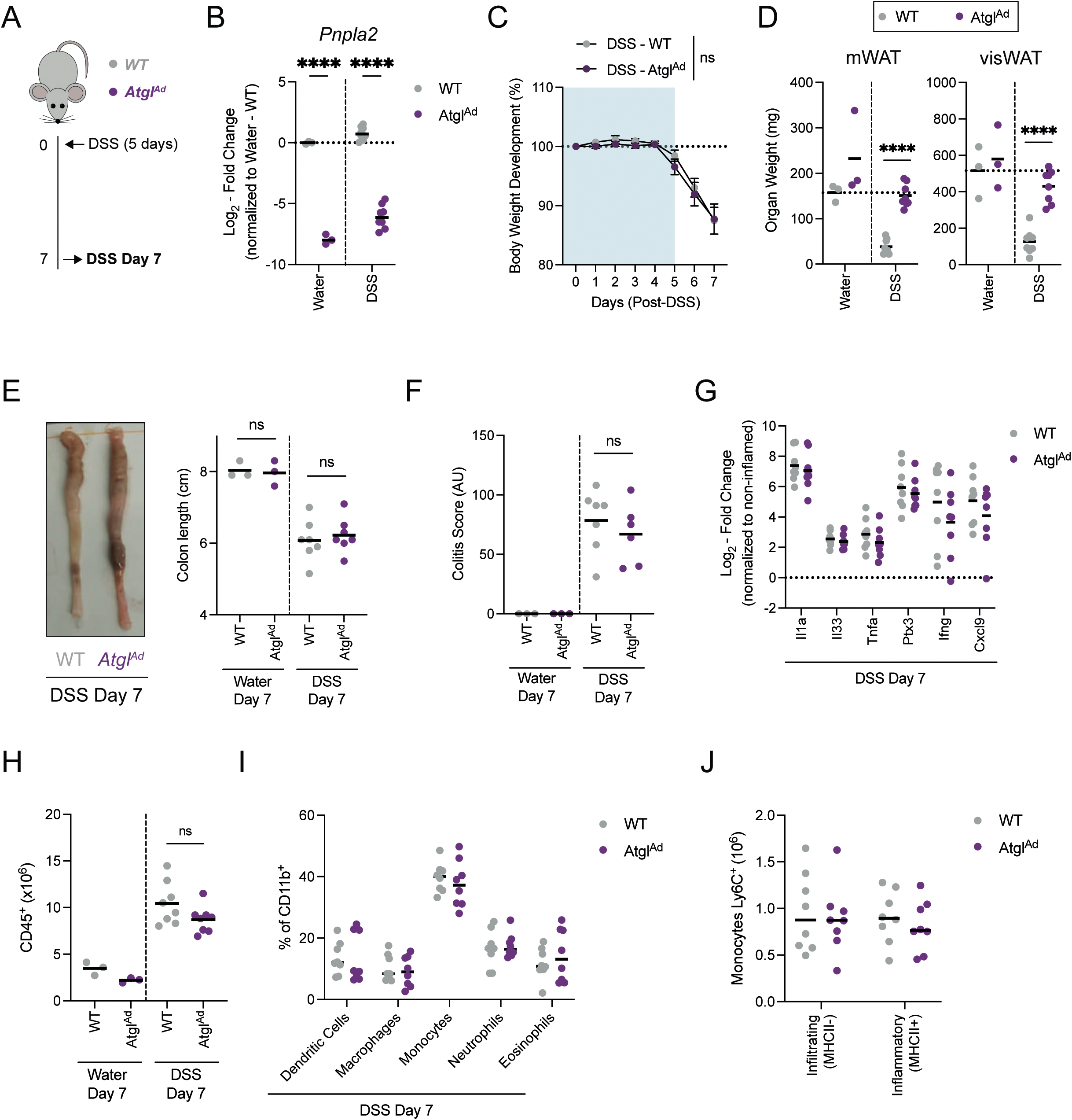
Adipocyte-specific loss of *Atgl* was dispensable for regulation of intestinal inflammation. (A) Schematic of experimental design. DSS-induced colitis was induced in sex-matched and age- matched littermates. (B) Representative quantification of knock-out efficiency measured on *Atgl* transcript level by qRT-PCR in purified primary visceral adipocyte (n = 3-8/group). (C) Body weight development upon DSS treatment (n = 3-8/group). (D) Tissue weights of mWAT and visWAT at day 7 after start of DSS (n = 3-8/group). (E) Colon length after DSS at day 7 (n = 3-8) pooled from two independent experiments. (F) Quantification of histological score at steady state (n = 3/group) and DSS colitis (n = 6-7/group). (G) Expression of pro-inflammatory cytokines in colon tissues at 7 days post-DSS induction (n = 8/group). Dotted line represents non-inflamed controls. (H) Absolute number CD45^+^ immune cells from colons at 7 days post-DSS induction (n = 3-8/group). (I) Frequency of myeloid cell population in colon at day 7 post-DSS induction (n = 8/group). (J) Absolute number of Ly6C^+^ monocytes discriminated by the absence or presence of MHCII for infiltrating and inflammatory monocytes respectively (n = 8/group). Data are represented as mean. (B,D-H,J) Unpaired Student’s t-test. (C,I) Two-Way ANOVA.

### Adipocyte autophagy loss promotes NRF2-mediated stress response and alters tissue oxylipin levels

To get a better understanding of pathways that may be affected by the loss of autophagy in adipocytes. We further analysed our transcriptomic data by splitting the dataset based on their condition and genotype. Reassuringly, visceral adipocytes from *Atg7^Ad^* mice had a strong reduction in *Atg7* levels and an increase in estrogen receptor 1 (*Esr1*) expression, due to the *Cre* transgene expression (Figure 6A- B). Across both treatment groups, we found a total of 32 genes being differentially regulated between WT and *Atg7^Ad^* visceral adipocytes. Six genes were differentially expressed under both water and DSS treatment conditions (Figure 6C). Due to the limited number of differentially expressed genes between *Atg7^Ad^* and wild-type adipocytes, we opted to look for altered pathway using ranked gene set enrichment analysis (GSEA)^31^, which includes genome-wide alterations to given gene sets of major cellular pathways. Upon DSS-induced colitis, we found that the xenobiotic pathway, was significantly enriched in *Atg7^Ad^* adipocytes (Figure 6D+S5A). Enzymes which are known for their role in xenobiotic metabolism such as the large family of cytochromes P450 monooxygenases and epoxide hydrolases (EPHX) metabolize and detoxify exogenous substrates and mediate the production of oxylipins from endogenous polyunsaturated fatty acids. The expression of many of the key genes involved in these processes are regulated by NRF2, a major transcription factor of the xenobiotic and oxidative stress responses. We found that *Ephx1* was consistently upregulated upon *Atg7* loss in adipocytes (Figure 6C). Remarkably, *Ephx1* expression was also increased in datasets obtained from other studies in which autophagy genes such as *Atg3* and *Beclin-1* were specifically deleted in adipocytes (Figure S5B- C)^19, 25^. Among the genes that were enriched in *Atg7*-deficient adipocytes were several other NRF2- target genes (Figure 6E). In agreement with an activation of the NRF2 pathway, NRF2 protein abundance was increased in *Atg7^Ad^* visceral adipose tissues (Figure 6F). Specificity for NRF2 activation was further confirmed since only NRF2 target gene *Ephx1* was transcriptionally upregulated, whereas *Ephx2* which is not controlled by NRF2 remained transcriptionally unchanged in autophagy-deficient adipocytes (Figure 6G). However, both EPHX1 and EPHX2 protein expression were increased in *Atg7^Ad^* adipose tissues (Figure 6H) suggesting that EPHX2 may be affected by autophagy deletion on a post- transcriptional level.

**Figure 6:**
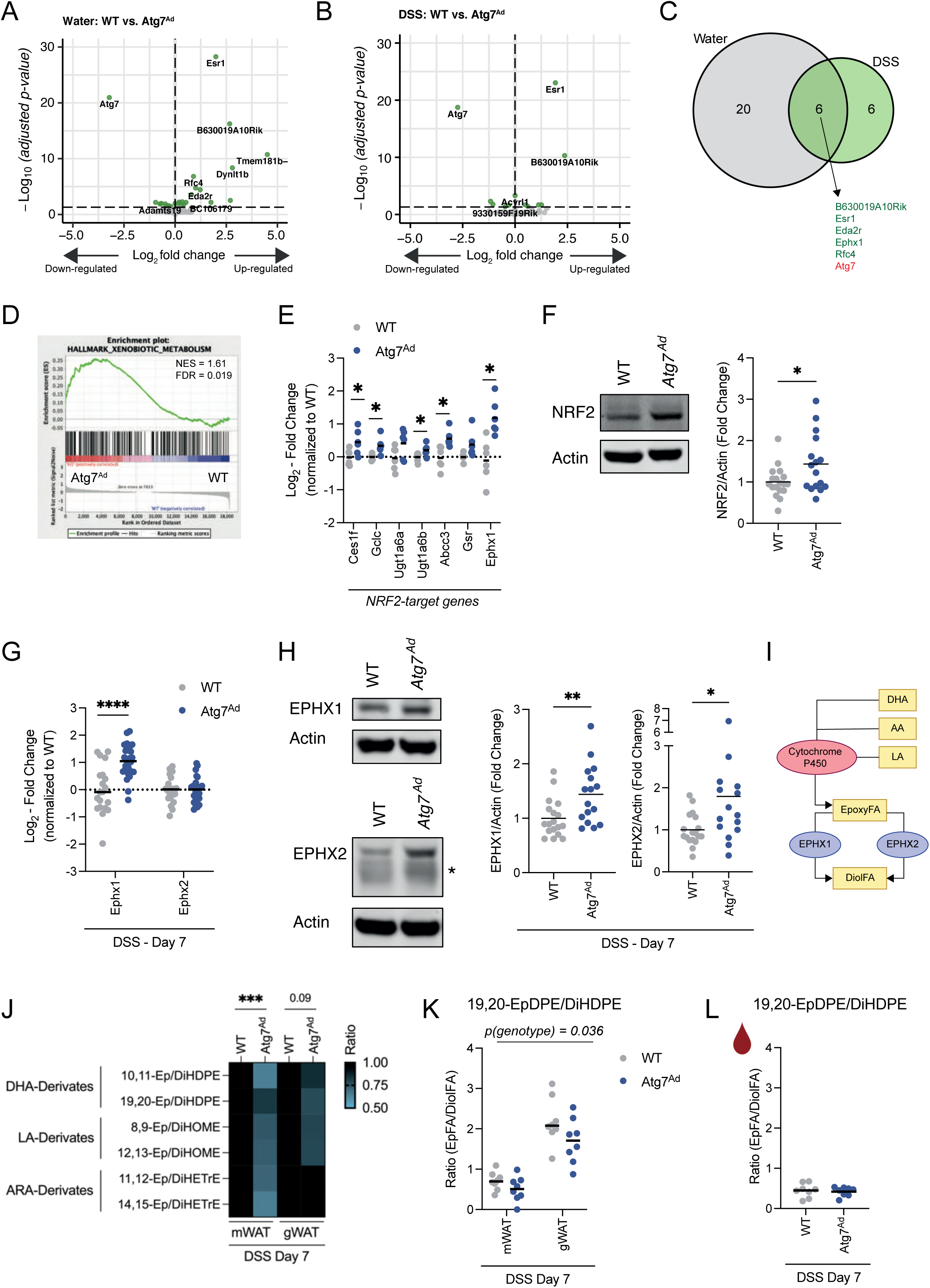
Adipocyte autophagy loss activates NRF2-EPHX1 pathway and alters intra-tissual oxylipin balance. (A) Differential gene expression in visceral adipocytes from water-treated WT and *Atg7^Ad^* animals two weeks after tamoxifen treatment. (B) Differential gene expression in visceral adipocytes from DSS-treated WT and *Atg7^Ad^* animals at day 7 post-DSS treatment. (C) Venn diagram of commonly regulated genes between *Atg7*-deficient and *Atg7*-sufficient adipocytes during water- or DSS-treatment. (D) GSEA enrichment analysis between *Atg7*-deficient and *Atg7*-sufficient adipocytes during DSS- treatment. (E) Fold change expression of NRF2-target genes in primary visceral adipocytes at day 7 after DSS induction from normalized counts of RNAseq dataset (n = 6/group). (F) Representative immunoblot for NRF2 protein expression and quantification (n = 16-18/group). (G) Transcriptional expression of *Ephx1 and Ephx2* in visceral adipocytes at day 7 after DSS induction pooled from three cohorts (n = 20-25/group). (H) Representative immunoblot of EPHX1 and EPHX2 in gonadal adipose tissues at day 7 after DSS induction. Asterix indicating non-specific band (n = 15-18/group). (I) Schematic overview of cytochrome P450-EPHX oxylipin pathway. (J) Normalized ratios of epoxy fatty acid to their corresponding diol fatty acids pairs in mWAT and gWAT (n = 6-8/group). (K) Ratio of 19,20-EpDPE:19,20-DiHDPE in mWAT and gWAT at day 7 after DSS induction (n = 8/group). (L) Ratio of 19,20-EpDPE:19,20-DiHDPE in blood plasma at day 7 after DSS induction (n =. 8/group). Data presented as mean. (E-H, L) Unpaired Student’s t-test. (J) One-Way ANOVA. (K) Two-Way ANOVA.

EPHX1, together with EPHX2, are central for the enzymatic conversion of cytochrome P450-derived oxylipins such as epoxy fatty acids (EpFA) to dihydroxy/diol fatty acids (DiolFA). EpFA have strong anti-inflammatory, analgesic and hypotensive activity, while DiolFA are less biologically active and are associated with more pro-inflammatory properties, respect (Figure 6I)^32^. In addition to the increase in EPHX1 and EPHX2 abundance in adipose tissues, we found that an enrichment of the oxylipin substrate docosahexaenoic acid (DHA) and a trend for arachidonic acid (ARA) in serum of *Atg7^Ad^* mice during DSS-induced inflammation (Figure S5D) suggesting a putative activation of the oxylipin pathway. We therefore tested whether the increased expression of EPHX enzymes would shift the balance of oxylipins in the tissue and, possibly, plasma. Indeed, loss of adipocyte autophagy alters the EpFA:DiolFA ratio towards a reduction of anti-inflammatory EpFAs in both mesentery and gonadal adipose tissues during DSS colitis (Figure 6J). This was consistently observed for all analyzed DHA-derived EpFAs which are important substrates for EPHX1 (Figure 6K)^15^. Strikingly, these effects appear to be locally restricted to the adipose tissues since no changes in these plasma oxylipin levels were observed (Figure 6L). In summary, these data suggest that loss of adipocyte autophagy activates NRF2 and increased the expression of EPHX enzymes promoting a local disbalance of EpFA:DiolFA. We hypothesize that this dysbalance in turn might alter the local inflammatory response in the adipose tissue to intestinal inflammation.

### Adipose tissues increase secretion of IL-10 in response to DSS-induced colitis in an autophagy-dependent manner

Since the changes in EpFA:DiolFA appeared to be locally restricted, we next wanted to identify which soluble factors may impact on gut inflammation. Evidence suggests that stimulation of macrophages with EpFA promotes the production of IL-10, while abundance of DiolFA can quench IL-10 production^33, 34^. Thus, we next tested whether cytokine production from the adipose tissue may contribute to systemic inflammation during DSS-induced colitis and assessed the secreted cytokine profile from mesenteric adipose tissues. We found that the mesenteric adipose tissue increases the secretion of several cytokines including the anti-inflammatory cytokine IL-10 in response to DSS (Figure 7A). We found that, upon DSS-induced colitis, F4/80^+^ macrophages are one of the major cell populations producing IL-10 in mesenteric and visceral adipose tissues, although mesenteric CD4^+^ T cells appear to contribute as well (Figure 7B). In adipose tissues of DSS treated mice, ATM frequencies were increased, but was comparable between the genotypes (Figure S7A). EpFA can alter macrophage polarization and increase M2-type macrophage marker expression such as CD206^35^. In line with reduced EpFA levels in the adipose tissue, *Atg7^Ad^* ATMs expressed reduced CD206 expression (Figure S7B). In addition, expression CD36, a lipid scavenging receptor which is commonly found on M2-type macrophages, was increased on the surface of ATMs in wild-type mice during DSS colitis and blunted in *Atg7^Ad^* mice (Figure S7C). Since ATMs are a prominent source of IL-10 production, we next tested whether IL-10 secretion from the mesenteric and gonadal adipose tissue was affected by adipocyte autophagy loss. Remarkably, disruption of adipocyte autophagy abolished DSS-induced IL-10 secretion from both mesenteric and gonadal adipose tissues, while having little impact on TNFα secretion, indicating that even adipose tissues that are not adjacent to the inflammation site contribute to the anti-inflammatory response (Figure 7C-D). Due to the important role of IL-10 in immune tolerance, we hypothesized that the reduction of IL-10 secretion from adipose tissues may translate into a systemic reduction of circulating IL-10 levels. Indeed, we found that while circulating IL-10 levels were significantly upregulated in DSS-treated wild-type mice compared to non-inflamed mice, their expression was diminished in *Atg7^Ad^* mice (Figure 7E). Taken together, the data suggest that, in line with a decreased availability of anti-inflammatory EpFA, adipose tissues from adipocyte autophagy-deficient mice have an impaired production and secretion of anti-inflammatory IL-10 to DSS induced colitis compared to wild-type mice.

**Figure 7:**
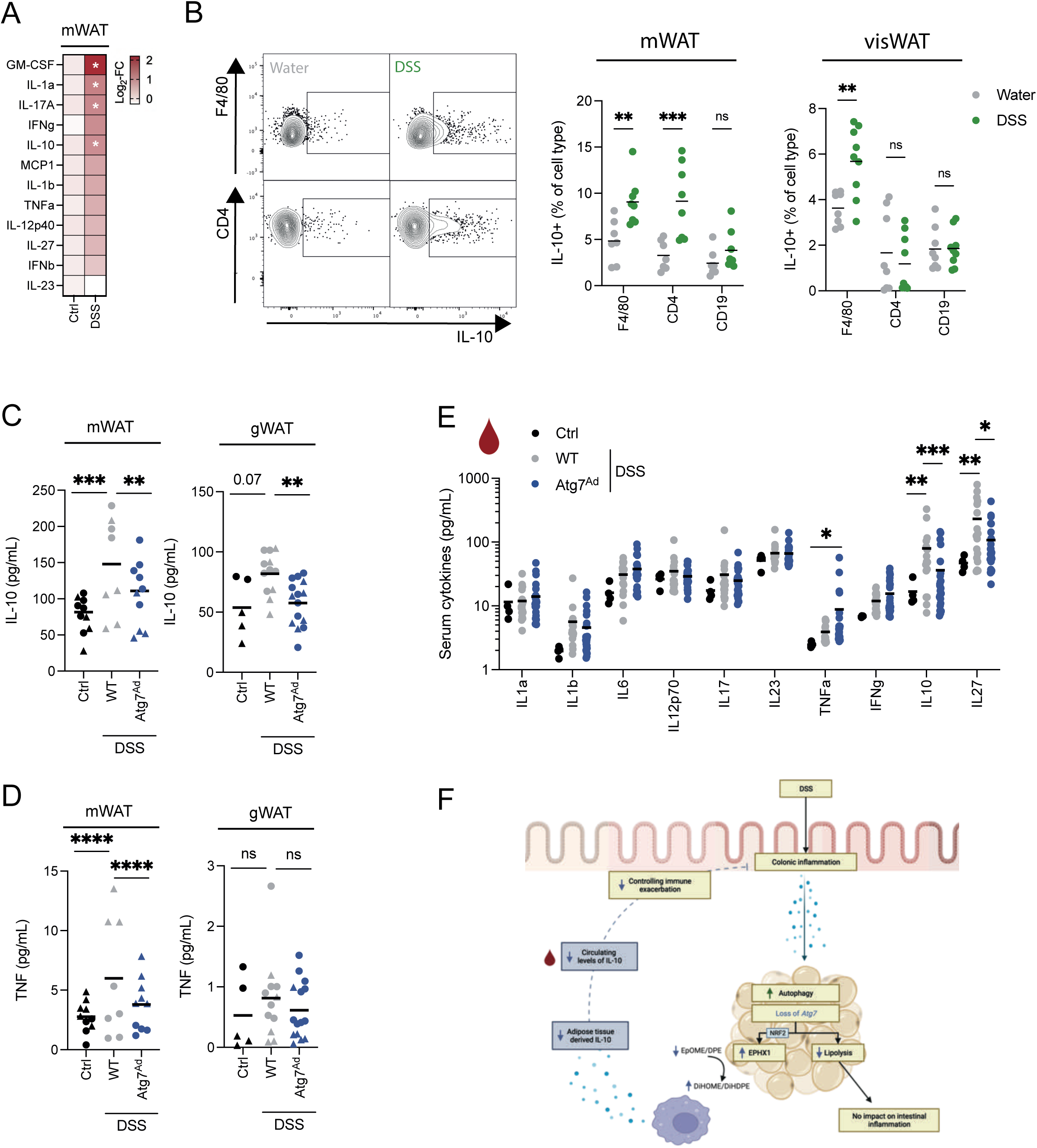
IL-10 is secreted from adipose tissues in an autophagy-dependent manner during DSS-induced colitis. (A) Colitis was induced in mice for 7 days and mesenteric adipose tissue explants were cultured with FBS. IL-10 secretion into the supernatant was measured after 24h of culture (n = 4-12). (B) Identification of IL-10-producing cells in adipose tissue upon DSS-induced colitis by flow cytometry. Representative FACS plots (left panel) and quantification from mesenteric (middle panel) and visceral adipose tissues (right panel) (n = 7-8/group). (C and D) Colitis was induced in mice for 7 days and adipose tissues were extracted and cultured for 6 hours in serum-starved medium. Secretion of (C) IL-10 and (D) TNFα from mesenteric (left panel) and gonadal adipose tissues (right panel) was measured by ELISA. Shapes identify individual experiments (n = 5-15/group). (E) Serum cytokines upon DSS-induced colitis at day 7 post-induction (n = 17-23/group) (F) Graphical summary of the anti-inflammatory fat-gut crosstalk during intestinal inflammation. Data are represented as mean. (A) One-Way ANOVA. (B,E) Two-Way ANOVA. (C,D) Two-Way ANOVA with regression for experiment.

## Discussion

Immune cells reside within distinct tissue environments, however, the impact of local metabolic cues on inflammatory processes remains incompletely understood^36^. Our results indicate that autophagy in mature adipocytes contributes to the balance of intra-tissual oxylipin levels. Further, we demonstrate that adipocytes autophagy is part of the anti-inflammatory immune response by promoting the release of IL-10 from adipose tissues. Autophagy-dependent secretion from adipose tissues contributes to systemic IL-10 levels, and limits inflammation at a distant tissue site, the colon. Therefore, our study provides novel insights into a cross-tissue anti-inflammatory mechanism, enabling the development of therapeutic approaches to target this crosstalk.

While polymorphisms in autophagy genes are well established as genetic risk factors for IBD, little is known about autophagy’s role in adipocytes in this disease. We found that autophagy is induced in visceral adipocytes upon DSS-induced colitis, which was marked, among others, by a transcriptional increase of *Atg8* homologues during peak inflammation. These observations parallel findings during muscle atrophy, where the expression of *Map1lc3b, Gabarapl1, Bnip3, Bnip3l* and *Vps34* is regulated via FOXO3 activation, which subsequently controls autophagy levels^37^. It appears plausible that a similar FOXO3-dependent mechanism occurs in adipocytes, especially since visceral adipocytes showed several transcriptomic and macroscopic changes reminiscent of a ‘cachexia-like’ phenotype. We hypothesize that intestinal-derived cues (such as TNFα or bacterial translocation) may promote systemic inflammation which is required for the induction of adipocyte autophagy and cachexia-like phenotype^38^.

Early studies found that autophagy is crucial for the normal differentiation of adipose tissues *in vivo*^18, 20^. However, the significance of autophagy in mature adipocytes remained unexplored until recently. Post-developmental ablation of autophagy in mature adipocytes decreased β-adrenergic receptor-induced lipolysis^19, 25^. Conversely, disruption of mTOR by genetic deletion of *Raptor* increases lipolytic output via autophagy^39^. It is likely that adipocyte autophagy controls lipolytic output via the degradation of key proteins involved in the lipolytic machinery such as described for perilipins in fibroblasts and adipocytes^40, 41^. We extended this knowledge by our finding that adipocyte autophagy also regulates TNFα-induced lipolysis and by this may fine-tune lipolytic output of adipocytes upon inflammatory stress conditions.

We further demonstrate that adipocyte lipolysis during DSS-induced colitis is driven predominantly through ATGL. Somewhat surprisingly, loss of adipocyte lipolysis had no impact on body weight loss or colonic inflammation, thus raising the question whether adipocyte lipolysis is beneficial or maladaptive in the context of this disease. These observations are reminiscent of findings during infection-associated cachexia, where deletion of the cytosolic lipases *Atgl* and *Hsl* had no impact on body weight loss^42^. In contrast, during cancer-associated cachexia, loss of these lipases prevents body weight loss suggesting that infection and inflammation models of cachexia act through distinct and yet to be identified biological pathways^43^.

Loss of adipocyte autophagy increased NRF2 stability, likely through the sequestration of its regulator KEAP1 as shown by Cai et al.^19^. Here we demonstrate for the first time that this antioxidant/xenobiotic pathway exacerbates an inflammatory disease. Increased expression of EPHX1 was paralleled by a dysbalance in oxylipins shifted towards decreased levels of EpFA and increased DiolFA. Similar to our findings, EPHX1 was recently found to convert in particularly omega-3 DHA substrates in adipocytes and liver^15, 44^. Since our data suggest a broader dysregulation of EpFA:DiolFA, it is likely that EPHX2, which was accumulated on protein level in *Atg7^Ad^* adipose tissues, may also contribute to the conversion of oxylipin substrates. Increasing evidence suggests that macrophages are regulated by oxylipins in their environment. Indeed, increased presence of omega-3 derived EpFA achieved either through inhibition of EPHX2 or through supplementation has been shown to promote CD206 expression and IL- 10 secretion^34, 35^. In line with the reduced presence of EpFA in *Atg7*-deficient adipose tissue, we found these two hallmarks of anti-inflammatory macrophages were equally decreased.

Importantly, this study underscores the importance of adipose-tissue derived IL-10 in controlling disease severity. Our findings of increased IL-10 secretion in visceral adipose tissues upon intestinal inflammation, confirmed findings from the Siegmund lab that mesenteric ATMs upregulate expression of IL-10 during intestinal inflammation in both human and mouse^45, 46^. In line with this, global loss of IL- 10 leads to exacerbation of intestinal inflammation^47^. The disruption in systemic IL-10 levels may also explain the reduced colonic expansion of FOXP3^+^ Tregs at resolution, since adequate IL-10 signalling is required for the expression of FOXP3 in intestinal Tregs^48^. Recent single cell transcriptomic analysis of immune cells resident in human creeping fat tissues revealed an anti-inflammatory and pro-repair role of ATMs, further supporting their beneficial role during intestinal inflammation^49^. Our data highlights how adipocyte dysfunction can impair this adipocyte-immune cell crosstalk suggesting that this communication may also exist in human pathology.

While ATMs accumulate in in creeping fat tissues of CD patients and in the mesentery of mice upon DSS-induced colitis^45^, it remains unclear how these macrophages are regulated during intestinal inflammation. We propose that oxylipins can shift macrophage polarization, in part, through their action as PPAR ligands^50^. which are important regulators of M2-type polarization and function^51, 52^. We found that the PPARγ-target gene CD36 is up-regulated during DSS-induced colitis on ATMs in wild-type mice and but not in ATMs from *Atg7^Ad^* mice. Similarly, the expression of CD36 on adipocytes can be controlled by oxylipin levels^53^. In addition, it is possible that oxylipin disbalance may also affect other IL-10 producing cell types in the adipose tissues such as Tregs which also rely on PPARγ for their accumulation and function^54^. The resulting reduction in systemic IL-10 levels prolongs pro-inflammatory programs at the distal inflammation site. As such, IL-10 signalling is required for intestinal macrophages to prevent pro-inflammatory exacerbation during DSS-induced colitis through inhibition of mTOR signalling which controls macrophage pro-inflammatory activity^47, 55^.

Overall, this study reveals that metabolically healthy adipose tissues are important regulators to prevent excessive inflammation during colitis. However, the function of adipose tissues in IBD may depend on the overall metabolic and disease state. The expansion of the mesentery during CD may initially be beneficial through prevention of bacterial translocation and signalling pathways poised to promote anti-inflammatory pathways, as shown here^45, 49^. However, sustained inflammation may ultimately subvert the function of the mesentery and lead to adipose tissue fibrosis and intestinal strictures^56^. Sustained tissue fibrosis results in tissue hypoxia^57^ and may impact on tissue oxylipin levels in creeping fat of CD patients.

Here, we demonstrate for the first time that adipocyte autophagy contributes to the intra-tissual balance of oxylipin levels and thus controls the anti-inflammatory immune response to intestinal tissue injury through regulation of adipose tissue derived IL-10 (as summarized in Figure 7F). It underlines the importance of local adipocyte-immune cell crosstalk through regulation of lipid mediators. This may present a broader local metabolic regulatory pathway to control immune responses to inflammation and infection.

## Methods

### Mice

*Adipoq-Cre^ERT2^* mice^58^ were purchased from Charles River, UK (JAX stock number: 025124) and were crossed to *Atg7* floxed mice^59^. Experimental cages were sex- and age-matched and balanced for genotypes. Genetic recombination was induced at 8-10 weeks of age by oral gavage of 4mg tamoxifen per mouse for five consecutive days. All experimental procedures were conducted two weeks after last tamoxifen administration. DSS-induced colitis was induced by 1.5-2% (w/v) DSS (MP Biomedicals, 160110) in drinking water. Mice were treated with DSS for five days and assessed at day 7, a peak inflammation time, or at day 14, a resolution time point. Constitutive *Adipoq-Cre x Pnpla2* floxed mice^60, 61^ (JAX stock number: 024278) were kindly provided by Prof. Rudolph Zechner. Wild-type C57BL/6J mice were purchased from Charles River, UK (JAX stock number: 0000664) or bred in-house. Mice were housed on a 12-hour dark/light cycle and fed *ad libitum*, under specific pathogen-free conditions. All animal experimentation was performed in accordance to approved procedures by the Local Review Committee and the Home Office under the project licence (PPL30/3388 and P01275425).

### Histopathology assessment

Distal, mid and proximal colon pieces were fixed in 10% neutral buffered formalin for 24 hours before washed and transferred into 70% ethanol. Tissue pieces from each sample were embedded in the same paraffin block and 5µm sections were subsequently stained with haematoxylin and eosin (H&E). Scoring of histology sections was executed in a blinded fashion according to a previously reported scoring system^21^. In brief, each section was assessed for the degree inflammation, the depth of tissue damage, possible crypt damages, with high scores signifying increased tissue damage. In addition, signs of regeneration (epithelial closure, crypt regeneration) were assessed, with high scores indicating delayed regeneration. Changes were multiplied with a factor classifying the involvement tissue area. Total score was calculated and presented.

### Adipose tissue and colon digestion

We collected mesenteric adipose tissue separate from a collective set of visceral adipose tissue depots (including omental, gonadal and retroperitoneal adipose tissue) to distinguish proximal versus distal effects of intestinal inflammation on adipose tissues. Adipose tissues were collected and digested in DMEM containing 1% fatty acid-free BSA (Sigma, 126609), 5% HEPES (Gibco, 15630-056), 0.2mg/mL Liberase TL (Roche, 5401020001) and 20µg/mL DNaseI (Roche, 11284932001). Tissues were minced in digestion medium and incubated for 25-30min at 37°C at 180rpm. Tissues were further broken down by pipetting using wide-bore tips and filtered through a 70µm mesh. Digestion was quenched by adding medium containing 2mM EDTA. Adipocyte and stromal vascular fraction were separated by centrifugation (700g, 10min) and collected for further downstream analysis.

Colon digestions were performed as previously described^62^. Colons were opened longitudinally and faecal content was removed by washing with PBS. Then colons were washed twice in RPMI containing 5% FBS and 5mM EDTA at 37°C under agitation. Tissues were minced and digested in RPMI supplemented with 5% FBS, 1mg/mL collagenase type VIII (Sigma) and 40μg/mL DNaseI (Roche). Cell suspension was strained through 40µm mesh and cells were subjected to downstream analysis.

### Flow Cytometry

Flow cytometry staining was performed as previously described^63^. Surface staining was performed by incubating cells with fluorochrome-conjugated antibodies (Biolegend, BD Bioscience, eBioscience) and LIVE/DEAD Fixable Stains (ThermoFischer) for 20min at 4°C. Cells were fixed with 4% PFA for 10min at room temperature. For intracellular staining of transcription factors, cells were fixed/permeabilized using the eBioscience™ Foxp3/ Transcription Factor Staining Set (00-5523-00, Invitrogen). For cytokine staining, cells were stimulated using Cell Activation cocktail (Biolegend) for 4h at 37°C in RPMI containing 10% FBS. After surface staining, cells were fixed and stained in Cytofix/CytoPerm (BD Bioscience) following manufacturer protocol. Samples were acquired on LSRII or Fortessa X-20 flow cytometers (BD Biosciences).

### Quantitative PCR

Adipocytes and adipose tissue RNA were extracted using TRI reagent (T9424, Sigma). Colon tissue RNA were extracted in RLT buffer containing 1,4-Dithiothreitol. Tissues were homogenised by lysis in 2mL tubes containing ceramic beads (KT03961-1-003.2, Bertin Instruments) using a Precellys 24 homogenizer (Bertin Instruments). RNA was purified following RNeasy Mini Kit (74104, Qiagen) manufacturer instructions. cDNA was synthesized following the High-Capacity RNA-to-cDNA™ kit protocol (4388950, ThermoFischer). Gene expression was assessed using validated TaqMan probes and run on a ViiA7 real-time PCR system. All data were collected by comparative Ct method either represented as relative expression (2^-ΔCt^) or fold change (2^-ΔΔCt^). Data were normalized to the two most stable housekeeping genes; for adipose tissues *Tbp* and *Rn18s* and for colon *Actb* and *Hprt*.

### Bulk RNA sequencing

Visceral adipocytes were isolated as floating fraction upon digestion. RNA was extracted and converted to cDNA as described above. PolyA libraries were prepared through end reparation, A-tailing and adapter ligation. Samples were then size-selected, multiplexed and sequenced using a NovaSeq6000. Raw read quality control was performed using pipeline readqc.py (https://github.com/cgat-developers/cgat-flow). Resulting reads were aligned to GRCm38/Mm10 reference genome using the pseudoalignment method kallisto^64^. Differential gene expression analysis was performed using DEseq2 v1.30.1^65^. Pathway enrichment analysis was performed on differentially expressed genes for “Biological Pathways” using clusterProfiler (v4.0) R package^66^. DESeq2 median of ratios were used for visualisation of expression levels. Heatmaps of selected gene sets were presented as z-scores using R package pheatmap. Gene enrichment analysis was performed using GSEA software using Hallmark gene sets^31^. R code is available under https://github.com/cleete/IBD-Adipocyte-Autophagy

### Lipolysis assays

Adipose tissues were collected and washed in PBS before subjected to lipolysis assays. For isoproterenol stimulation, adipose tissues were cut into small tissue pieces and incubated in serum-free DMEM - High Glucose (Sigma, D5796) with 2% fatty acid-free BSA (Sigma, 126579) in the absence or presence of 10µM isoproterenol (Sigma, I6504) for the indicated time. TNFα-induced lipolysis was induced as previously described^41^. In brief, small adipose tissue pieces were cultured in DMEM – High Glucose for 24 hours in the absence or presence of 100ng/mL recombinant TNFα (Peprotech, 315- 01A) and then transferred into serum-free DMEM containing 2% fatty acid free BSA for 3 hours. Supernatants were collected and FFA concentration normalized to adipose tissue input.

### Adipose tissue explant cultures

Gonadal or mesenteric adipose tissue explants were collected from mice at indicated time points. For autophagic flux measurements, explants (∼50-100mg) were cultured for DMEM supplemented with 10% FBS (Sigma, F9665) and 100U/ml Pen-Strep for 4h in the absence or presence of lysosomal inhibitors 100nM Bafilomycin A1 and 20mM ammonium chloride. Explants were washed in PBS before collection and then frozen at -80°C until proteins were extracted for immunoblotting. For measurement of cytokine secretion, adipose tissue explants were cultured for 6h in DMEM/High Modified (D6429, Sigma) with 100U/ml Pen-Strep in the absence of FBS. Supernatant was collected, spun down (400g, 5min) to remove cell debris and then frozen until further analysis.

### Free fatty acid analysis

Total supernatant and serum FFA levels were measured using Free Fatty Acid Assay Quantification Kit (ab65341, Abcam). For detailed analysis of FFA species, lipids were extracted by Folch’s method ^67^ and subsequently run on a one-dimensional thin layer chromatography (TLC) using a 10x10cm silica gel G plate in a hexane/diethyl ether/acetic acid (80:20:1, by vol.) solvent system. Separated FFA were used for fatty acid methyl esters (FAMEs) preparation through addition of 2.5% H2SO4 solution in dry methanol/toluene (2:1 (v/v)) at 70°C for 2h. A known amount of C17:0 was added as an internal standard for quantification. FAMEs were extracted with HPLC grade hexane. A Clarus 500 gas chromatograph with a flame ionizing detector (FID) (Perkin-Elmer) and fitted with a 30m x 0.25mm i.d. capillary column (Elite 225, Perkin Elmer) was used for separation and analysis of FAs. The oven temperature was programmed as follows: 170°C for 3min, increased to 220°C at 4°C/min), and then held at 220°C for 15min. FAMEs were identified routinely by comparing retention times of peaks with those of G411 FA standards (Nu-Chek Prep Inc). TotalChrom software (Perkin-Elmer) was used for data acquisition and quantification.

### Oxylipin analysis

Oxylipins were analyzed by means of liquid chromatography mass spectrometry^68, 69^. The plasma samples were analyzed following protein precipitation and solid-phase extraction on reversed phase/anion exchange cartridges^68, 69^. The adipose tissue was homogenized in a ball mill and oxylipins were extracted with a mixture of chloroform and *iso*-propanol following solid-phase extraction on an amino propyl SPE cartridge^70, 71^. Oxylipin concentrations were determined by external calibration with internal standards^68, 69^.

### Immunoblotting

Autophagic flux in adipose tissues was measured by incubating adipose tissue explants from experimental animals in RPMI in the absence or presence of lysosomal inhibitors 100nM Bafilomycin A1 and 20mM NH4Cl for 4 hours. DMSO was used as ‘vehicle’ control. Adipose tissues were collected and snap frozen. Protein extraction was performed as previously described^72^. In brief, 500µL of lysis buffer containing protease inhibitors (04693159001, Roche) and phosphoStop (04906837001, Roche) were added per 100mg of tissue. Cells were lysed using Qiagen TissueLyser II. Tissues were incubated on ice for 1h and lipid contamination was removed via serial centrifugation and transfer of internatant into fresh tubes. Protein concentration was determined by BCA Protein Assay Kit (23227, Thermo Scientific). Total of 15-30µg protein were separated on a 4-12% Bis-Tris SDS PAGE and transferred using BioRad Turbo Blot (1704156, BioRad) onto PVDF membrane. Membranes were blocked in TBST containing 5% milk. Primary antibodies were used at indicated concentration overnight. Membranes were visualized using IRDye secondary antibodies (LICOR). Band quantification of Western Blots was performed on ImageJ. Autophagic flux was calculated as: (LC3-II (Inh) – LC3-II (Veh))/(LC3-II (Veh)), as previously described^73^.

### Transmission electron microscopy

Mice were sacrificed by increasing concentrations of CO2. Adipose tissues were excised, cut into small 1-2mm pieces and immediately fixed in pre-warmed (37 °C) primary fixative containing 2.5% glutaraldehyde and 4% formaldehyde in 0.1M sodium cacodylate buffer, pH7.2 for 2 hours at room temperature and then stored in the fixative at 4 °C until further processing. Samples were then washed for 2x 45 min in 0.1M sodium cacodylate buffer (pH 7.2) at room temperature with rotation, transferred to carrier baskets and processed for EM using a Leica AMW automated microwave processing unit. Briefly, this included three washes with 0.1M sodium cacodylate buffer, pH 7.2, one wash with 50mM glycine in 0.1M sodium cacodylate buffer to quench free aldehydes, secondary fixation with 1% osmium tetroxide + 1.5% potassium ferricyanide in 0.1M sodium cacodylate buffer, six water washes, tertiary fixation with 2% uranyl acetate, two water washes, then dehydration with ethanol from 30%, 50%, 70%, 90%, 95% to 100% (repeated twice). All of these steps were performed at 37 °C and 15-20W for 1-2 mins each, with the exception of the osmium and uranyl acetate steps, which were for 12 min and 9 min respectively. Samples were infiltrated with TAAB Hard Plus epoxy resin to 100% resin in the AMW and then processed manually at room temperature for the remaining steps. Samples were transferred to 2ml tubes filled with fresh resin, centrifuged for ∼2mins at 2000g (to help improve resin infiltration), then incubated at room temperature overnight with rotation. The following day, the resin was removed and replaced with fresh resin, then the samples were centrifuged as above and incubated at room temperature with rotation for ∼3 hrs. This step was repeated and then tissue pieces were transferred to individual Beem capsules filled with fresh resin and polymerised for 48 hrs at 60 °C. Once polymerised, blocks were sectioned using a Diatome diamond knife on a Leica UC7 Ultramicrotome. Ultrathin (90nm) sections were transferred onto 200 mesh copper grids and then post-stained with lead citrate for 5 mins, washed and air dried. Grids were imaged with a Thermo Fisher Tecnai 12 TEM (operated at 120 kV) using a Gatan OneView camera.

### Extracellular cytokine measurements

Serum samples were collected by cardiac puncture and collected in Microtainer tubes (365978, BD Bioscience). Samples were centrifuged for 90sec at 15,000g and serum aliquots were snap-frozen until further analysis. Global inflammatory cytokine analysis of supernatants of adipose tissue explant cultures and serum were performed using LEGENDPlex™ Mouse Inflammation Panel (740446, Biolegend). TNFα and IL-10 levels were measured by TNFα Mouse Uncoated ELISA Kit (88-7324-86, Invitrogen) and IL-10 Mouse Uncoated ELISA Kit (88-7105-86, Invitrogen), respectively. Adipose tissue derived cytokine levels were normalized to input tissue weight.

### Statistical Analysis

Data were tested for normality before applying parametric or non-parametric testing. For two normally-distributed groups, unpaired Student’s tests were applied. Comparisons across more than two experimental groups were performed using One-Way or Two-Way ANOVA with Šídák multiple testing correction. Data were considered statistically significant when p<0.05 (*p<0.05, **p<0.01, ***p<0.001, ****p<0.0001). Typically, data were pooled from at least two experiments, if not otherwise indicated, and presented as mean. Data was visualized and statistics calculated in either GraphPad Prism 9 or R software.

## Acknowledgements

We thank Patricia Cotta Moreira, Daniel Andrew and Mino Medghalchi from the Kennedy Institute animal facility for their excellent care and assistance of animal well-being. Dr. Nicholas Illot, Dr. Alina Janney and Dr. Luca Baù for their help with R coding. Discussions about experimental design for the transcriptomic experiments and sequencing were performed with help of Prof. Stephen Samson, Dr. Moustafa Attar and the Oxford Genomics Centre. Histology was performed with the help from the Kennedy Institute Histology Facility, especially Dr. Ida Parisi. This work was supported by grants from the Wellcome Trust (Investigator award 103830/Z/14/Z and 220784/Z/20/Z to A.K.S., Investigator award 212240/Z/18/Z to F.P., PhD studentship award 203803/Z16/Z to F.C.R., PhD studentship award 108869/Z/15/Z to S.K.W.), the Kenneth Rainin Foundation (Innovator award 20210017 to A.K.S. and F.P., jointly), the Kennedy Trust Studentship (KEN192001 to K.P.), the Marie Sklodowska-Curie - European Fellowship (893676 to M.B.) Blood Cancer UK (15026 and 17012 to C.M.E). Li-cor Odyssey imager was funded by ERC AdG 670930.

## Authors Contribution

Conceptualization, F.C.R, M.F and K.A.S.; Methodology, F.C.R, M.F., N.K., I.G., E.J., N.H.S.; Formal Analysis, F.C.R., M.F. N.K.; Investigation, F.C.R., M.F., N.K., N.H.S., M.P., G.A., I.G., S.K.W., E.J., M.B., K.P., P.H.; Writing – Original Draft, F.C.R., A.K.S; Writing – Review and Editing, M.F., M.P., N.K. N.H.S., G.A., I.G., S.K.W., E.J., M.B., K.P., P.H., H.S.S., C.M.E., A.K.S.; Visualization, F.C.R, Supervision, M.F., H.S.S., C.M.E, F.P., N.H.S., A.K.S.; Funding Acquisition, F.C.R., F.P., A.K.S.

## Declaration of Interests

F.P. received research support or consultancy fees from Roche, Janssen, GSK, Novartis and Genentech. A,K.S. received consultancy fees from Calico, Oxford Healthspan, The Longevity Lab.

---

**Supplementary Figure 1: DSS leads to efficient induction of intestinal inflammation.**

(A) Representative H&E staining of colon histology and quantification at day 7 after DSS colitis induction (n = 3/group) from one independent experiment.
(B) Colon length measured after 1.5-2% DSS colitis regime at day 7 (n = 6-7/group).
(C) Spleen weight and mesenteric lymph node weight after 1.5% colitis regime at day 7 (n = 9/group).
(D) TNFα levels in serum were measured in wild-type mice at day 7 after water and DSS treatment (n=5/group).
(E) Absolute number of colonic CD45^+^ immune cells at day 7 post-DSS treatment (n = 6-7/group).
(F) Frequency of CD11b^+^ myeloid cells, CD3^+^ T cells and CD19^+^ B cells in colon at day 7 post-DSS treatment (n = 6-7/group).

Data are represented as mean. (A-E) Unpaired Student’s t-test.

**Supplementary Figure 2: Expansion of intestinal Treg populations is blunted in adipocyte autophagy-deficient mice without affecting intestinal resolution.**

(A) Schematic of experimental design. Sex-matched and age-matched littermates were treated with DSS for five days and mice were sacrificed 14 days after start of DSS treatment.
(B) Colon length from non-inflamed control mice (n = 8/group), adipocyte autophagy-sufficient WT mice and adipocyte autophagy-deficient mice (n = 12/group).
(C) Representative H&E staining images (10x magnification) of distal colon sections and quantification of histopathological score; n=7-13 pooled from two independent experiment.
(D) Frequency (left panel) and absolute number (right panel) of CD4^+^ FOXP3^+^ cells in the colon at day 14 post-DSS treatment (n = 8-11).
(E) Frequency of peripheral and thymic Treg (pTreg and tTreg, respectively) cell populations in colon at day 14 post-DSS treatment (n = 6-11).

Data are represented as mean. (B-D) One-Way ANOVA. (E) Two-Way ANOVA.

**Supplementary Figure 3: Primary visceral adipocytes are enriched in macroautophagy pathway genes.**

(A) Heatmap representing differentially expressed genes associated with macroautophagy during DSS- induced colitis in visceral adipocytes.
(B) Normalized counts of selected key enzymes and proteins involved in the lipolysis pathway in visceral adipocytes (n = 12/group).

Data are represented as mean. (B) Unpaired Student’s t-test.

**Supplementary Figure 4: Loss of adipocyte autophagy had no effects on adipose tissue wasting and circulating levels of leptin and adiponectin.**

(A) Adipose tissue mass at steady state or at day 7 post-DSS induction (n = 7-9/group).
(B) Circulating levels of adiponectin (n = 3-12/group).
(C) Circulating levels of leptin (n = 4-12/group).

Data are represented as mean. (A) Unpaired Student’s t-test. (B,C) One-Way ANOVA.

**Supplementary Figure 5: Loss of autophagy-related genes results in the induction of epoxy hydrolases in adipocytes.**

(A) GSEA enrichment analysis between *Atg7*-deficient and *Atg7*-sufficient adipocytes during DSS- treatment.
(B) Fragments per kilobase of exon per million mapped fragments (FPKM) counts from bulk RNAseq dataset of Cai et al.^19^ (n = 4/group)
(C) Fragments per kilobase of exon per million mapped fragments (FPKM) counts from bulk RNAseq dataset of Son et al.^25^ (n = 3/group).
(D) Frequency of cytochrome P450 substrates for oxylipin production in serum (n =. 13-20/group).

Data are represented as mean. (B-D) Unpaired Student’s t-test.

**Supplementary Figure 6: Adipocyte autophagy loss results in shifts of macrophage polarization.**

(A) Frequency of adipose tissue macrophages among all immune cells in adipose tissues (n =. 7- 8/group).
(B) Expression of CD206 on adipose tissue macrophages in visceral adipose tissues (n = 6-12/group).
(C) Expression of CD36 on adipose tissue macrophages. Dotted line represents uninflamed controls (n = 5-7/group).

Data are represented as mean. (A-C) Unpaired Student’s t-test.

## References

1. Clarke, A. J. & Simon, A. K. Autophagy in the renewal, differentiation and homeostasis of immune cells. Nat Rev Immunol 19, 170–183, doi:10.1038/s41577-018-0095-2 (2019).

2. Deretic, V. Autophagy in inflammation, infection, and immunometabolism. Immunity 54, 437–453, doi:10.1016/j.immuni.2021.01.018 (2021).

3. Klionsky, D. J. et al. Autophagy in major human diseases. EMBO J 40, e108863, doi:10.15252/embj.2021108863 (2021).

4. Hampe, J. et al. A genome-wide association scan of nonsynonymous SNPs identifies a susceptibility variant for Crohn disease in ATG16L1. Nat Genet 39, 207–211, doi:10.1038/ng1954 (2007).

5. Jostins, L. et al. Host-microbe interactions have shaped the genetic architecture of inflammatory bowel disease. Nature 491, 119–124, doi:10.1038/nature11582 (2012).

6. McCarroll, S. A. et al. Deletion polymorphism upstream of IRGM associated with altered IRGM expression and Crohn’s disease. Nat Genet 40, 1107–1112, doi:10.1038/ng.215 (2008).

7. Cadwell, K. et al. A key role for autophagy and the autophagy gene Atg16l1 in mouse and human intestinal Paneth cells. Nature 456, 259–263, doi:10.1038/nature07416 (2008).

8. Cadwell, K., Patel, K. K., Komatsu, M., Virgin, H. W. t. & Stappenbeck, T. S. A common role for Atg16L1, Atg5 and Atg7 in small intestinal Paneth cells and Crohn disease. Autophagy 5, 250–252, doi:10.4161/auto.5.2.7560 (2009).

9. Kabat, A. M. et al. The autophagy gene Atg16l1 differentially regulates Treg and TH2 cells to control intestinal inflammation. Elife 5, e12444, doi:10.7554/eLife.12444 (2016).

10. Sheehan, A. L., Warren, B. F., Gear, M. W. & Shepherd, N. A. Fat-wrapping in Crohn’s disease: pathological basis and relevance to surgical practice. Br J Surg 79, 955–958, doi:10.1002/bjs.1800790934 (1992).

11. Trim, W. V. & Lynch, L. Immune and non-immune functions of adipose tissue leukocytes. Nat Rev Immunol, doi:10.1038/s41577-021-00635-7 (2021).

12. Russo, L. & Lumeng, C. N. Properties and functions of adipose tissue macrophages in obesity. Immunology 155, 407–417, doi:10.1111/imm.13002 (2018).

13. Huang, S. C. et al. Cell-intrinsic lysosomal lipolysis is essential for alternative activation of macrophages. Nat Immunol 15, 846–855, doi:10.1038/ni.2956 (2014).

14. Klein-Wieringa, I. R. et al. Adipocytes modulate the phenotype of human macrophages through secreted lipids. J Immunol 191, 1356–1363, doi:10.4049/jimmunol.1203074 (2013).

15. Edin, M. L. et al. Epoxide hydrolase 1 (EPHX1) hydrolyzes epoxyeicosanoids and impairs cardiac recovery after ischemia. J Biol Chem 293, 3281–3292, doi:10.1074/jbc.RA117.000298 (2018).

16. Gilroy, D. W. et al. CYP450-derived oxylipins mediate inflammatory resolution. Proc Natl Acad Sci U S A 113, E3240–3249, doi:10.1073/pnas.1521453113 (2016).

17. Imig, J. D. & Hammock, B. D. Soluble epoxide hydrolase as a therapeutic target for cardiovascular diseases. Nat Rev Drug Discov 8, 794–805, doi:10.1038/nrd2875 (2009).

18. Singh, R. et al. Autophagy regulates adipose mass and differentiation in mice. J Clin Invest 119, 3329–3339, doi:10.1172/JCI39228 (2009).

19. Cai, J. et al. Autophagy Ablation in Adipocytes Induces Insulin Resistance and Reveals Roles for Lipid Peroxide and Nrf2 Signaling in Adipose-Liver Crosstalk. Cell Rep 25, 1708–1717 e1705, doi:10.1016/j.celrep.2018.10.040 (2018).

20. Zhang, Y. et al. Adipose-specific deletion of autophagy-related gene 7 (atg7) in mice reveals a role in adipogenesis. Proc Natl Acad Sci U S A 106, 19860–19865, doi:10.1073/pnas.0906048106 (2009).

21. Dieleman, L. A. et al. Chronic experimental colitis induced by dextran sulphate sodium (DSS) is characterized by Th1 and Th2 cytokines. Clin Exp Immunol 114, 385–391, doi:10.1046/j.1365-2249.1998.00728.x (1998).

22. Whibley, N., Tucci, A. & Powrie, F. Regulatory T cell adaptation in the intestine and skin. Nat Immunol 20, 386–396, doi:10.1038/s41590-019-0351-z (2019).

23. Oliva, M. et al. The impact of sex on gene expression across human tissues. Science 369, doi:10.1126/science.aba3066 (2020).

24. Baazim, H., Antonio-Herrera, L. & Bergthaler, A. The interplay of immunology and cachexia in infection and cancer. Nat Rev Immunol, doi:10.1038/s41577-021-00624-w (2021).

25. Son, Y. et al. Adipocyte-specific Beclin1 deletion impairs lipolysis and mitochondrial integrity in adipose tissue. Mol Metab 39, 101005, doi:10.1016/j.molmet.2020.101005 (2020).

26. Friedrich, M., Pohin, M. & Powrie, F. Cytokine Networks in the Pathophysiology of Inflammatory Bowel Disease. Immunity 50, 992–1006, doi:10.1016/j.immuni.2019.03.017 (2019).

27. Cawthorn, W. P. & Sethi, J. K. TNF-alpha and adipocyte biology. FEBS Lett 582, 117–131, doi:10.1016/j.febslet.2007.11.051 (2008).

28. Siegmund, B., Lehr, H. A. & Fantuzzi, G. Leptin: a pivotal mediator of intestinal inflammation in mice. Gastroenterology 122, 2011–2025, doi:10.1053/gast.2002.33631 (2002).

29. Weidinger, C., Ziegler, J. F., Letizia, M., Schmidt, F. & Siegmund, B. Adipokines and Their Role in Intestinal Inflammation. Front Immunol 9, 1974, doi:10.3389/fimmu.2018.01974 (2018).

30. Schweiger, M. et al. Adipose triglyceride lipase and hormone-sensitive lipase are the major enzymes in adipose tissue triacylglycerol catabolism. J Biol Chem 281, 40236–40241, doi:10.1074/jbc.M608048200 (2006).

31. Subramanian, A. et al. Gene set enrichment analysis: a knowledge-based approach for interpreting genome-wide expression profiles. Proc Natl Acad Sci U S A 102, 15545–15550, doi:10.1073/pnas.0506580102 (2005).

32. McReynolds, C., Morisseau, C., Wagner, K. & Hammock, B. Epoxy Fatty Acids Are Promising Targets for Treatment of Pain, Cardiovascular Disease and Other Indications Characterized by Mitochondrial Dysfunction, Endoplasmic Stress and Inflammation. Adv Exp Med Biol 1274, 71–99, doi:10.1007/978-3-030-50621-6_5 (2020).

33. Levan, S. R. et al. Elevated faecal 12,13-diHOME concentration in neonates at high risk for asthma is produced by gut bacteria and impedes immune tolerance. Nat Microbiol 4, 1851–1861, doi:10.1038/s41564-019-0498-2 (2019).

34. McDougle, D. R. et al. Anti-inflammatory omega-3 endocannabinoid epoxides. Proc Natl Acad Sci U S A 114, E6034–E6043, doi:10.1073/pnas.1610325114 (2017).

35. Lopez-Vicario, C. et al. Inhibition of soluble epoxide hydrolase modulates inflammation and autophagy in obese adipose tissue and liver: role for omega-3 epoxides. Proc Natl Acad Sci U S A 112, 536–541, doi:10.1073/pnas.1422590112 (2015).

36. Richter, F. C., Obba, S. & Simon, A. K. Local exchange of metabolites shapes immunity. Immunology 155, 309–319, doi:10.1111/imm.12978 (2018).

37. Mammucari, C. et al. FoxO3 controls autophagy in skeletal muscle in vivo. Cell Metab 6, 458–471, doi:10.1016/j.cmet.2007.11.001 (2007).

38. Rivera, E. D., Coffey, J. C., Walsh, D. & Ehrenpreis, E. D. The Mesentery, Systemic Inflammation, and Crohn’s Disease. Inflamm Bowel Dis 25, 226–234, doi:10.1093/ibd/izy201 (2019).

39. Zhang, X. et al. Sustained activation of autophagy suppresses adipocyte maturation via a lipolysis-dependent mechanism. Autophagy 16, 1668–1682, doi:10.1080/15548627.2019.1703355 (2020).

40. Kaushik, S. & Cuervo, A. M. AMPK-dependent phosphorylation of lipid droplet protein PLIN2 triggers its degradation by CMA. Autophagy 12, 432–438, doi:10.1080/15548627.2015.1124226 (2016).

41. Ju, L. et al. Obesity-associated inflammation triggers an autophagy-lysosomal response in adipocytes and causes degradation of perilipin 1. Cell Death Dis 10, 121, doi:10.1038/s41419-019-1393-8 (2019).

42. Baazim, H. et al. CD8(+) T cells induce cachexia during chronic viral infection. Nat Immunol 20, 701–710, doi:10.1038/s41590-019-0397-y (2019).

43. Das, S. K. et al. Adipose triglyceride lipase contributes to cancer-associated cachexia. Science 333, 233–238, doi:10.1126/science.1198973 (2011).

44. Gautheron, J. et al. EPHX1 mutations cause a lipoatrophic diabetes syndrome due to impaired epoxide hydrolysis and increased cellular senescence. Elife 10, doi:10.7554/eLife.68445 (2021).

45. Batra, A. et al. Mesenteric fat - control site for bacterial translocation in colitis? Mucosal Immunol 5, 580–591, doi:10.1038/mi.2012.33 (2012).

46. Kredel, L. I. et al. Adipokines from local fat cells shape the macrophage compartment of the creeping fat in Crohn’s disease. Gut 62, 852–862, doi:10.1136/gutjnl-2011-301424 (2013).

47. Li, B., Alli, R., Vogel, P. & Geiger, T. L. IL-10 modulates DSS-induced colitis through a macrophage-ROS-NO axis. Mucosal Immunol 7, 869–878, doi:10.1038/mi.2013.103 (2014).

48. Murai, M. et al. Interleukin 10 acts on regulatory T cells to maintain expression of the transcription factor Foxp3 and suppressive function in mice with colitis. Nat Immunol 10, 1178–1184, doi:10.1038/ni.1791 (2009).

49. Ha, C. W. Y. et al. Translocation of Viable Gut Microbiota to Mesenteric Adipose Drives Formation of Creeping Fat in Humans. Cell 183, 666–683 e617, doi:10.1016/j.cell.2020.09.009 (2020).

50. Overby, H. et al. Soluble Epoxide Hydrolase Inhibition by t-TUCB Promotes Brown Adipogenesis and Reduces Serum Triglycerides in Diet-Induced Obesity. Int J Mol Sci 21, doi:10.3390/ijms21197039 (2020).

51. Odegaard, J. I. et al. Alternative M2 activation of Kupffer cells by PPARdelta ameliorates obesity-induced insulin resistance. Cell Metab 7, 496–507, doi:10.1016/j.cmet.2008.04.003 (2008).

52. Odegaard, J. I. et al. Macrophage-specific PPARgamma controls alternative activation and improves insulin resistance. Nature 447, 1116–1120, doi:10.1038/nature05894 (2007).

53. Lynes, M. D. et al. The cold-induced lipokine 12,13-diHOME promotes fatty acid transport into brown adipose tissue. Nat Med 23, 631–637, doi:10.1038/nm.4297 (2017).

54. Cipolletta, D. et al. PPAR-gamma is a major driver of the accumulation and phenotype of adipose tissue Treg cells. Nature 486, 549–553, doi:10.1038/nature11132 (2012).

55. Ip, W. K. E., Hoshi, N., Shouval, D. S., Snapper, S. & Medzhitov, R. Anti-inflammatory effect of IL-10 mediated by metabolic reprogramming of macrophages. Science 356, 513–519, doi:10.1126/science.aal3535 (2017).

56. Mao, R. et al. The Mesenteric Fat and Intestinal Muscle Interface: Creeping Fat Influencing Stricture Formation in Crohn’s Disease. Inflamm Bowel Dis 25, 421–426, doi:10.1093/ibd/izy331 (2019).

57. Zuo, L. et al. Mesenteric Adipocyte Dysfunction in Crohn’s Disease is Associated with Hypoxia. Inflamm Bowel Dis 22, 114–126, doi:10.1097/MIB.0000000000000571 (2016).

58. Sassmann, A., Offermanns, S. & Wettschureck, N. Tamoxifen-inducible Cre-mediated recombination in adipocytes. Genesis 48, 618–625, doi:10.1002/dvg.20665 (2010).

59. Komatsu, M. et al. Impairment of starvation-induced and constitutive autophagy in Atg7- deficient mice. J Cell Biol 169, 425–434, doi:10.1083/jcb.200412022 (2005).

60. Schoiswohl, G. et al. Impact of Reduced ATGL-Mediated Adipocyte Lipolysis on Obesity-Associated Insulin Resistance and Inflammation in Male Mice. Endocrinology 156, 3610–3624, doi:10.1210/en.2015-1322 (2015).

61. Sitnick, M. T. et al. Skeletal muscle triacylglycerol hydrolysis does not influence metabolic complications of obesity. Diabetes 62, 3350–3361, doi:10.2337/db13-0500 (2013).

62. Danne, C. et al. A Large Polysaccharide Produced by Helicobacter hepaticus Induces an Anti-inflammatory Gene Signature in Macrophages. Cell Host Microbe 22, 733–745 e735, doi:10.1016/j.chom.2017.11.002 (2017).

63. Riffelmacher, T. et al. Autophagy-Dependent Generation of Free Fatty Acids Is Critical for Normal Neutrophil Differentiation. Immunity 47, 466–480 e465, doi:10.1016/j.immuni.2017.08.005 (2017).

64. Bray, N. L., Pimentel, H., Melsted, P. & Pachter, L. Near-optimal probabilistic RNA-seq quantification. Nat Biotechnol 34, 525–527, doi:10.1038/nbt.3519 (2016).

65. Love, M. I., Huber, W. & Anders, S. Moderated estimation of fold change and dispersion for RNA-seq data with DESeq2. Genome Biol 15, 550, doi:10.1186/s13059-014-0550-8 (2014).

66. Wu, T. et al. clusterProfiler 4.0: A universal enrichment tool for interpreting omics data. Innovation (N Y) 2, 100141, doi:10.1016/j.xinn.2021.100141 (2021).

67. Folch, J., Lees, M. & Sloane Stanley, G. H. A simple method for the isolation and purification of total lipides from animal tissues. J Biol Chem 226, 497–509 (1957).

68. Rund, K. M. et al. Development of an LC-ESI(-)-MS/MS method for the simultaneous quantification of 35 isoprostanes and isofurans derived from the major n3- and n6-PUFAs. Anal Chim Acta 1037, 63–74, doi:10.1016/j.aca.2017.11.002 (2018).

69. Kutzner, L. et al. Development of an Optimized LC-MS Method for the Detection of Specialized Pro-Resolving Mediators in Biological Samples. Front Pharmacol 10, 169, doi:10.3389/fphar.2019.00169 (2019).

70. Koch, E., Wiebel, M., Hopmann, C., Kampschulte, N. & Schebb, N. H. Rapid quantification of fatty acids in plant oils and biological samples by LC-MS. Anal Bioanal Chem 413, 5439–5451, doi:10.1007/s00216-021-03525-y (2021).

71. Koch, E., Kampschulte, N. & Schebb, N. H. Comprehensive Analysis of Fatty Acid and Oxylipin Patterns in n3-PUFA Supplements. J Agric Food Chem 70, 3979–3988, doi:10.1021/acs.jafc.1c07743 (2022).

72. An, Y. A. & Scherer, P. E. Mouse Adipose Tissue Protein Extraction. Bio Protoc 10, e3631, doi:10.21769/BioProtoc.3631 (2020).

73. Zhang, H. et al. Polyamines Control eIF5A Hypusination, TFEB Translation, and Autophagy to Reverse B Cell Senescence. Mol Cell 76, 110–125 e119, doi:10.1016/j.molcel.2019.08.005 (2019).

