## Supplementary Figures for "Adipocyte autophagy limits gut inflammation by controlling oxylipin levels"

### Supplementary Figure 1

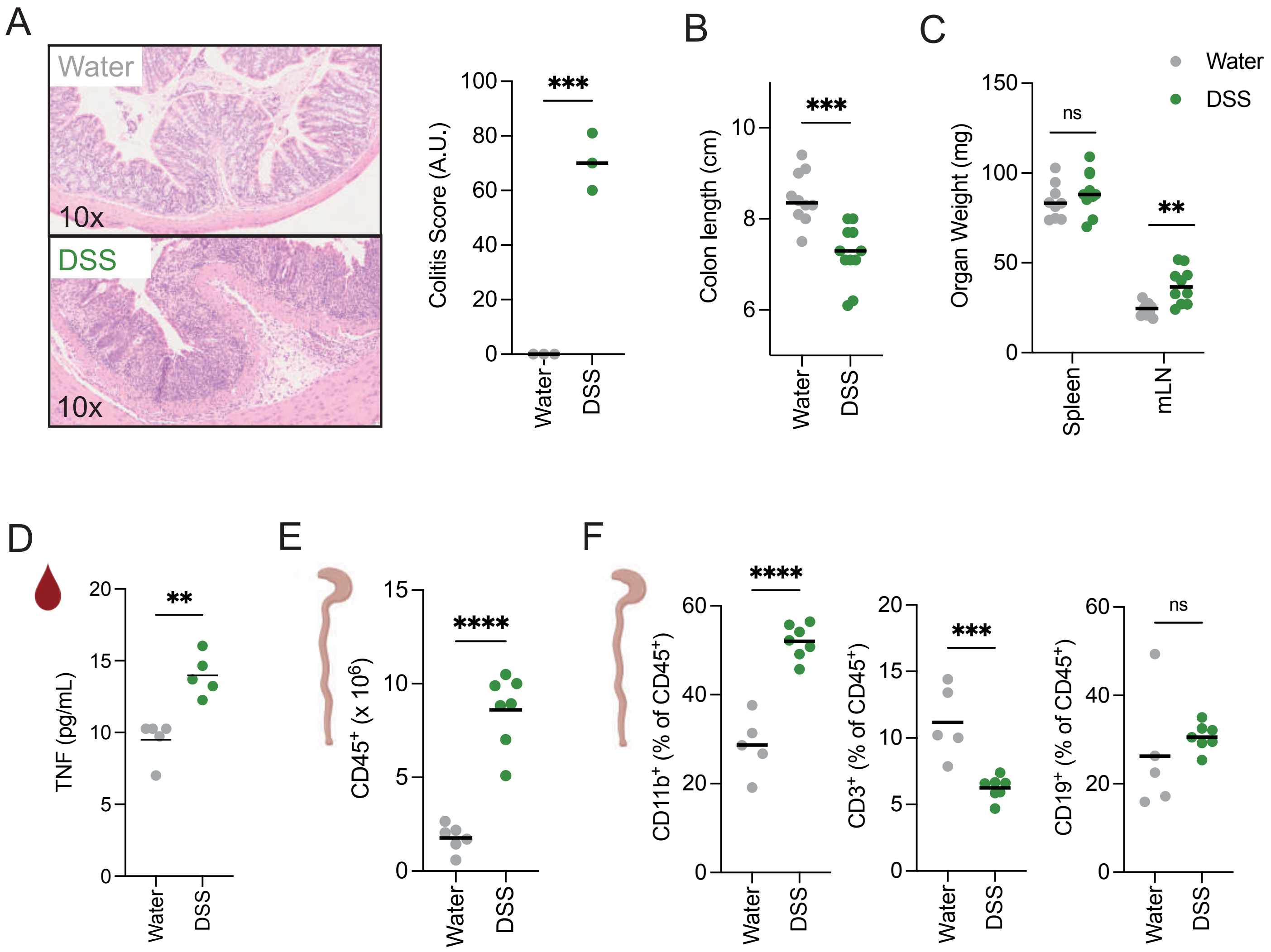

### Supplementary Figure 2

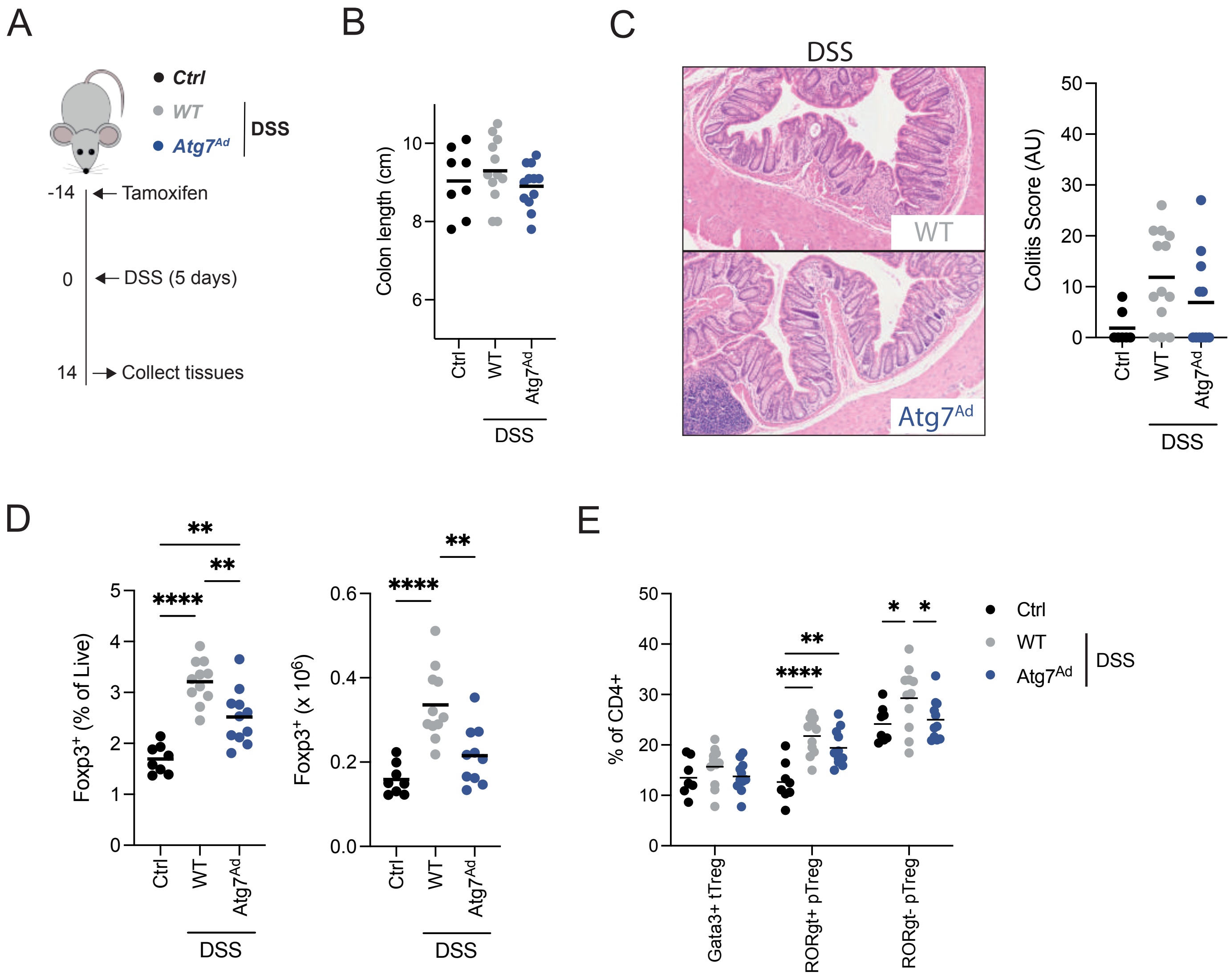

### Supplementary Figure 3

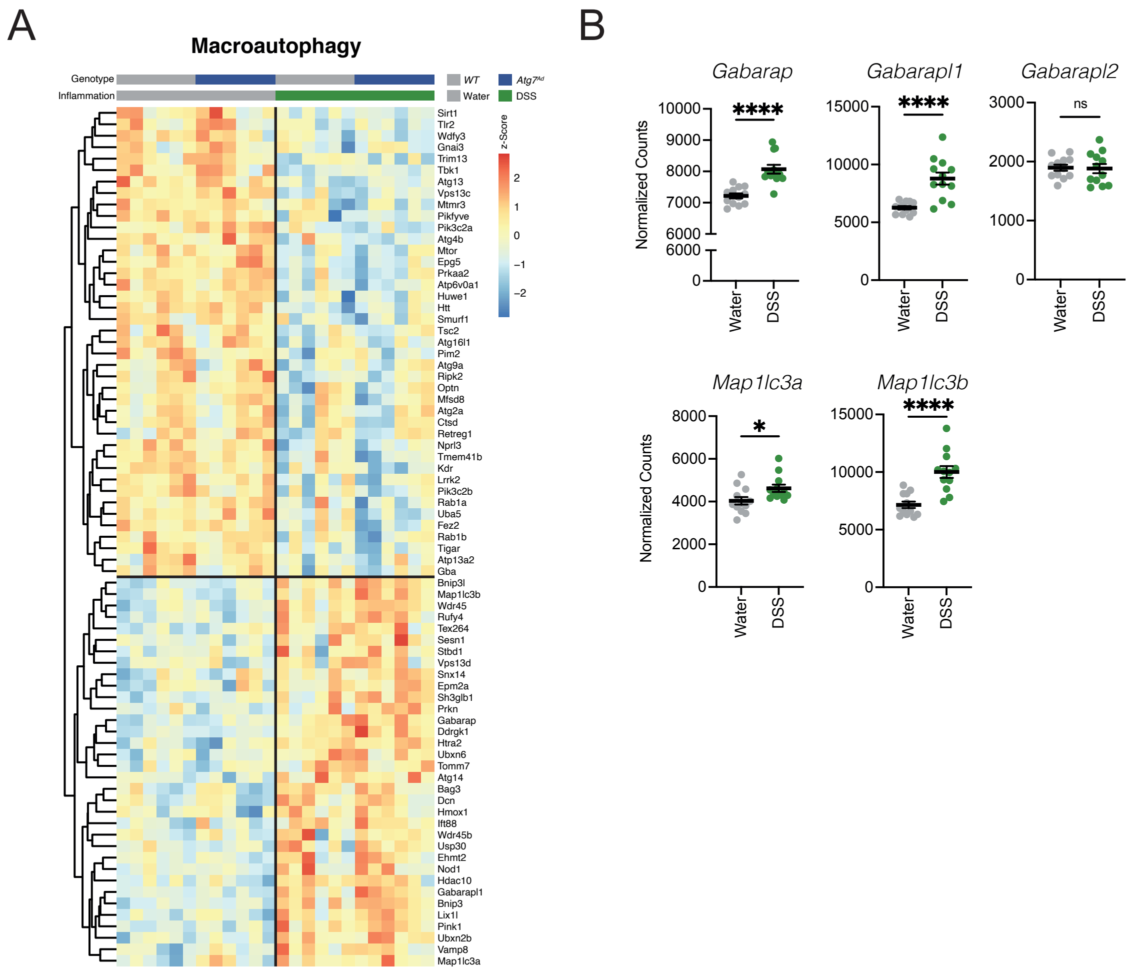

### Supplementary Figure 4

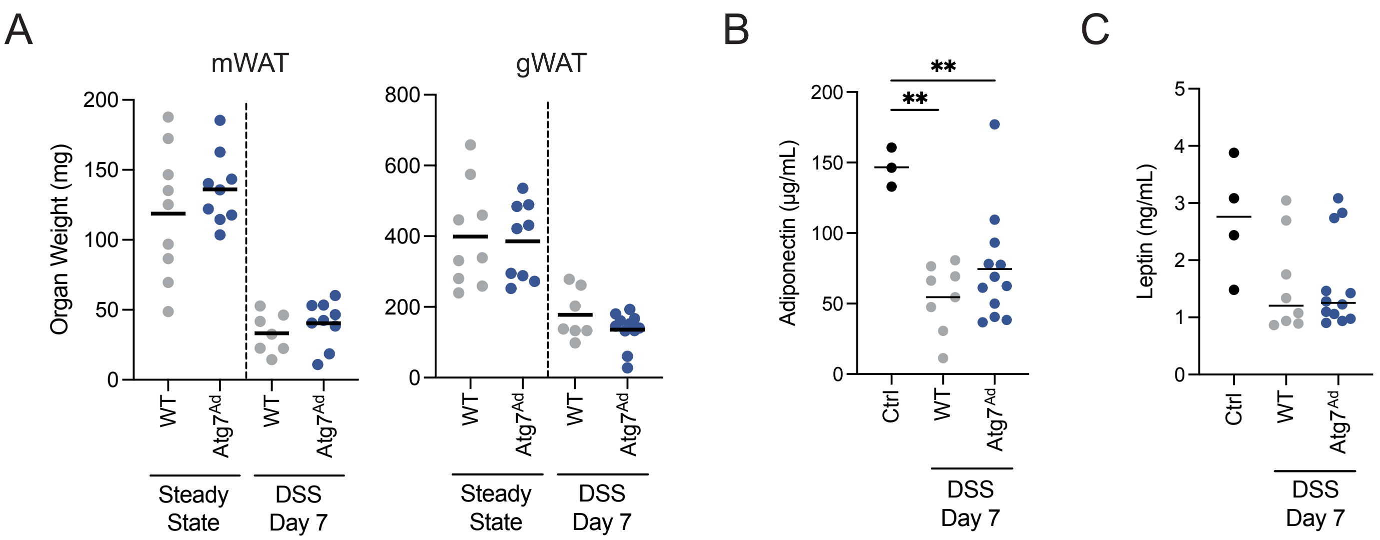

### Supplementary Figure 5

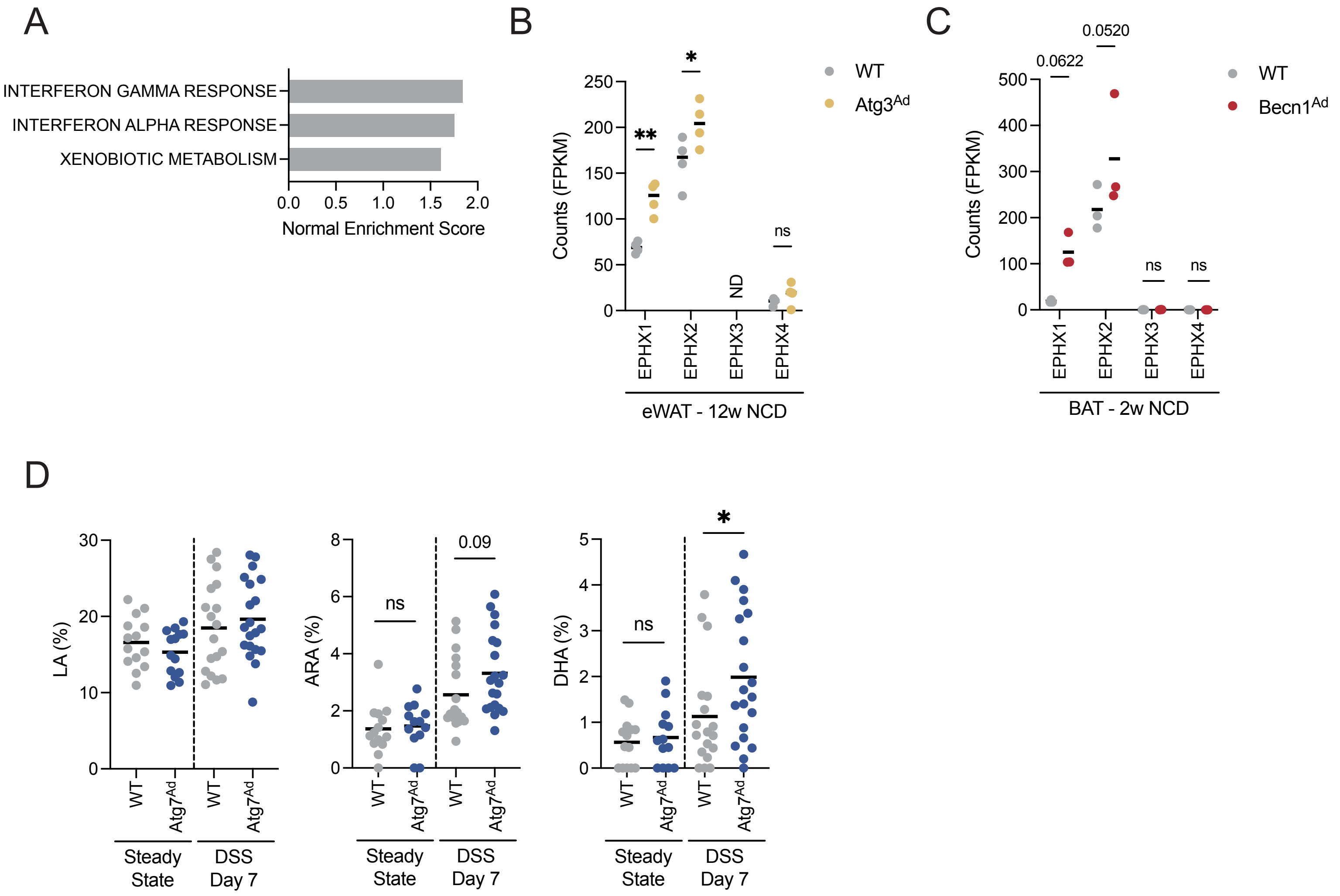

### Supplementary Figure 6

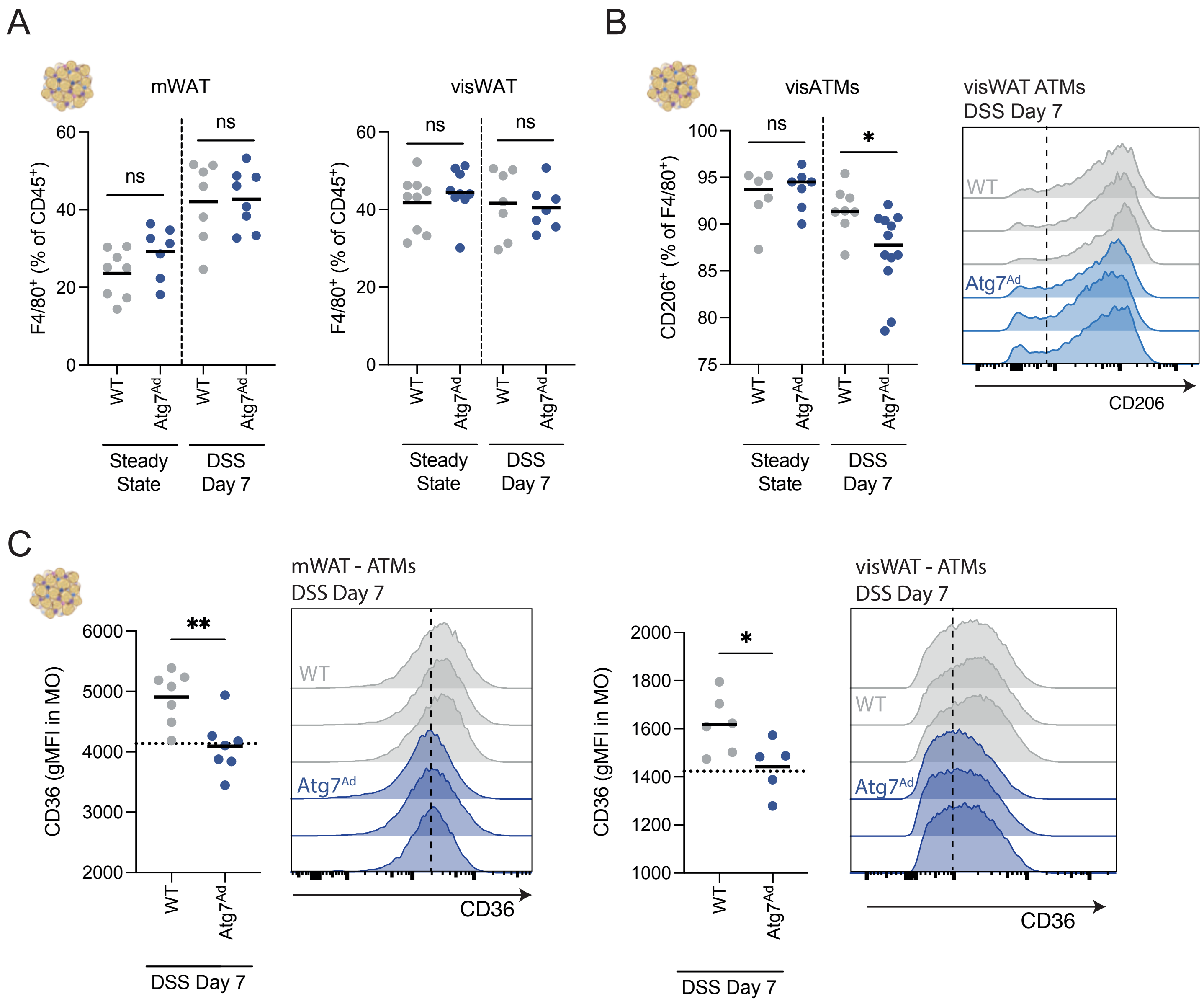
